# Leveraging the dominant-negative effect of the kuru-protective G127V prion protein variant as a novel therapeutic strategy

**DOI:** 10.64898/2026.02.17.703887

**Authors:** Jean R.P. Gatdula, Isabel C. Orbe, Samantha G. Tolton, Linnea M. Saunders, Janelle S. Vultaggio, Robert C.C. Mercer, Jason C. Bartz, Glenn C. Telling, Hasier Eraña, Joaquín Castilla, David A. Harris

## Abstract

Prion diseases are fatal neurodegenerative disorders with no approved therapies that halt or reverse disease progression. Given that cellular prion protein (PrP^C^) expression is required for prion propagation and neurotoxicity, reducing its expression is a promising therapeutic strategy. However, complete PrP ablation, as seen in knockout models, causes subtle developmental and behavioral abnormalities, raising concerns about long-term safety. Here, we explore a complementary strategy that harnesses the dominant-negative effect of the naturally protective G127V PrP variant found in kuru-resistant individuals in Papua New Guinea. In CAD5 cell lines, we demonstrate that inducible expression of G126V PrP (the mouse equivalent of human G127V) along with WT PrP prevents and suppresses prion infection in a dose-dependent manner. Extending this approach to CAD5 cells that express bank vole PrP, we further show that the protective effect of G127V spans a wide range of naturally and artificially derived prion strains, highlighting the generality of the dominant-negative approach. Remarkably, prion resistance persists even after G126V expression had ceased, indicating a sustained protective effect that could obviate the need for continuous transgene expression in a therapeutic setting. Finally, we find that anchorless, recombinant G127V PrP retains a potent dominant-negative activity, suggesting the use of this protein as a biological therapeutic. Together, these findings define a framework for development of G127V, a naturally protective and evolutionarily selected PrP variant, as a therapeutic agent to treat or prevent prion diseases.

## INTRODUCTION

Prion diseases are a group of rare, fatal, and transmissible neurodegenerative disorders that affect both humans and animals (Prusiner, 1982, Prusiner, 1998). These diseases are characterized by the misfolding of the non-infectious, cellular form of the prion protein (PrP^C^) into its infectious, β-sheet rich conformer (PrP^Sc^), which accumulates in the brain and drives neuronal dysfunction, degeneration, and death (Prusiner et al., 1981). Despite decades of investigation, there are currently no approved therapeutics that halt or reverse prion disease progression. Therapeutic efforts to date have largely focused on three strategies: (1) preventing PrP^C^ conversion to PrP^Sc^ using small molecules, ligands, or antibodies (Yamaguchi et al., 2019, Nicoll et al., 2010, Masone et al., 2023, White et al., 2003, Mead et al., 2022, Mercer et al., 2024, Mercer and Harris, 2019, Trevitt and Collinge, 2006); (2) reducing the synthesis of PrP^C^, using antisense oligonucleotides (ASOs), siRNAs, or epigenetic methods (Chou et al., 2025, Minikel et al., 2020); and (3) accelerating the degradation or clearance of PrP^Sc^ using lysosomal activity enhancers or other means (Mercer et al., 2025). Each of these approaches has strengths and weaknesses, and all are subject to the limitation of delivery of the therapeutic agent to the central nervous system (CNS).

In this study, we explore an alternative therapeutic strategy for preventing PrP^Sc^ formation without reducing or eliminating PrP expression, and instead takes advantage of a naturally occurring polymorphism in the *PRNP* gene. The G127V polymorphism was first identified in individuals of the Fore linguistic group in Papua New Guinea who were protected against kuru, a prion disease that was endemic in this group due to their practice of endocannibalism (Mead et al., 2009). Previous studies in humanized transgenic mice have shown that the G127V variant in a homozygous state confers complete protection against multiple CJD prion strains and kuru (Asante et al., 2015). Structurally, the G127V substitution locks PrP into a conformation that is conversion-incompetent, providing a molecular explanation for its remarkable protective effect. In addition to its ability to resist misfolding, the G127V variant acts as a potent, dominant-negative inhibitor of prion propagation by co-expressed wild-type PrP in transgenic mice (Asante et al., 2015).

In a previous study, we took advantage of the properties of G127V PrP to determine that PrP^Sc^ molecules generated on the neuronal cell membrane are the proximate triggers for prion synaptotoxic signaling (Gatdula et al., 2026). Here, we leverage this evolutionarily selected, protective mutation to block the propagation of PrP^Sc^ in cultured cell models, providing the framework for its development into an effective, broadly applicable therapeutic modality. We demonstrate the capacity of G126V PrP (the mouse homologue of human G127V) to prevent *de novo* prion infection and to suppress established infection caused by a range of natural and artificial prion strains in a dose-dependent manner. We show that, remarkably, this protective effect is sustained, even after expression of the variant protein is terminated, raising the possibility that it produces a long-lasting alteration in prion propagation. Finally, we establish that a soluble, recombinant form of the variant protein also has protective properties, laying the groundwork for its use as a biological therapeutic.

## METHODS

### Mammalian cell culture

Parental CAD5 and CAD5.Prnp^-/-^ cells were kindly provided by Joel Watts (Univ. of Toronto). CAD5 cells were maintained in Complete Opti-MEM Reduced Serum Medium (Thermo Fisher #31985088) supplemented with 10% (v/v) fetal bovine serum (FBS) and 1% (v/v) penicillin/streptomycin (Thermo Fisher #15140122). Cells were maintained in a humidified incubator containing 5% CO2 at 37 °C. Cells were passaged every 3 or 4 days at a dilution of 1:5 using 0.25% trypsin-EDTA (Fisher Scientific, MT25053CI). Stable cell lines were maintained using 200 µg/mL of G418 or 2 µg/mL of Puromycin.

### Constructs

The G126V (mouse) or G127V (bank vole) mutations were introduced into pcDNA3.1 (+)-Puro vector by restriction digest using BamHI and XbaI. WT bvPrP (I109) or G127V bvPrP (I109) inserts containing BamHI and XbaI restriction sites on the 5’ and 3’ ends, respectively, were generated using gBlocks from Integrated DNA Technologies (IDT). WT bvPrP (I109) in a pcDNA3-G418 vector (from Joel Watts) was used for stable co-expression experiments.

Dox inducible plasmids were made by introducing G126V PrP-IRES2-eGFP or WT PrP-IRES2-eGFP inserts containing BamHI and NheI restriction sites on the 5’ and 3’ ends into a Tet-On 3G Inducible Expression System vector (TakaraBio, #631168) using restriction digest and ligation. These inserts were generated using gBlocks from IDT.

Recombinant bvPrP plasmids were made by introducing codon-optimized WT bvPrP (I109) and G127V bvPrP (I109) inserts containing NheI and XhoI restriction sites on the 5’ and 3’ ends, into a pET41 plasmid (from Byron Caughey) using restriction digest and ligation. Inserts were generated using gBlocks from IDT.

### Generation of stably transfected or transduced CAD5 cells lines expressing PrP constructs

Parental CAD5 or CAD5.Prnp^-/-^ cells were transfected using Lipofectamine 3000 (Invitrogen # L3000001) according to the manufacturer’s directions. Murine PrP (WT or G126V) constructs were in a pcDNA3.1-Puro backbone plasmid while bank vole PrP (WT or G127V) constructs were in either a pcDNA3.1-Puro or a pcDNA3-G418 backbone plasmid. Following transfection, cells were selected over 6 or 10 days in medium containing 2 µg/ml of puromycin or 500 µg/ml of G418. The cells are then pooled and expanded without sub-cloning.

For transduction experiments, wild-type CAD5 or CAD5.Prnp^-/-^ cells were transduced using complete Opti-MEM media containing live lentiviruses. Briefly, old media was replaced with lentiviral media containing 2 µg/mL of Polybrene for better transduction efficiency. The cells are then spinoculated by centrifuging the plated cells for 1 hr. at 500x g for 30 minutes at 37 °C. The lentiviral medium was then replaced with fresh complete Opti-MEM media, after which cells recovered for 24 hours in the incubator. Cells were then selected over 6 days in medium containing 2 µg/ml of puromycin. The cells are then pooled and expanded without sub-cloning.

### Lentiviral production

Production of lentiviruses was performed as described previously (Gatdula et al., 2026). Lenti-X HEK293T cells were plated on a gelatin-coated dish and cultured in complete Opti-MEM. After 24 hours, cells were transfected with the packaging plasmids (psPAX2 and pMD2.G) and the transfer plasmid using polyethylenimine (PEI) at a transfer:psPAX2:pMD2.G ratio of 2:1:0.8. After 24 hours, the medium was replaced. Lentivirus-containing medium was collected after 24 hours and 48 hours post-media change. The lentiviral media was then pooled and centrifuged for 900x g for 5 min, and the supernatant was then filtered through a 0.45 μm filter. The filtered media is then aliquoted and stored in -80 °C.

### Prion-infected brain homogenates

Prion infected brains (mouse or hamster) were collected from C57BL/6 *Prnp^+/+^* mice inoculated with RML, 22L or ME7 prions, or from inoculated, C57BL/6 *Prnp^+/+^*control mice. Brains were homogenized using two rounds of 30 seconds of shaking with silica homogenizer beads. Stocks of brain homogenates were prepared at a final concentration of 10% in 1x PBS and stored in -80 °C. Prion infection of cell lines was carried out by applying 0.1% or 1% of brain homogenate in cell culture medium for 24 hours.

### CWD-infected cell homogenates

Cell homogenates were prepared as described (Bourkas et al., 2019) from confluent dishes of RK13 cells expressing either deer PrP or elk PrP that were chronically infected with Chronic Wasting Disease (CWD) prions from North American deer (designated 5E9 cells) or elk (designated 7F4 cells), respectively (a generous gift from Glenn Telling at Colorado State Univ.) (Bian et al., 2010). Cells were scraped into PBS, and then homogenized for 30s using a bead beater homogenizer followed by incubation on ice for 5 min. This was done three times. Cell homogenates are incubated with benzonase at a concentration of 50 units/mL for 30 min at 37°C, followed by centrifugation at 100x g to remove debris. Homogenates are stored at -80 °C. Prion infection of cell lines was carried out by applying cell homogenates (100 µg of protein in culture medium) for 24 hours.

### Recombinant PrP^Sc^ infection of cell lines

The three recombinant PrP^Sc^ preparations used here (from Joaquín Castilla and Hasier Eraña), which display distinct properties when inoculated into mice, were generated spontaneously (unseeded misfolding) by Protein Misfolding Shaking Amplification (PMSA) as described previously for other murine recombinant prions (Pérez-Castro et al., 2025). Briefly, PMSA substrate containing recombinant WT mouse PrP, complemented with dextran sulfate and glass beads was subjected to serial 24 h rounds of PMSA until first detection of PK-resistant PrP.

Replicates with different electrophoretic migration profiles after PK digestion were selected and their infectious nature and specific strain features were confirmed through intracerebral inoculation in WT mice. They are designated by the providers as stML-01-Dx, btML-89-Dx, and btML-90-Dx, and are referred to here as preparations A, B, and C, respectively, in Fig. 4G. Prion infection of cell lines was carried out by applying 100 ng of recombinant PrP^Sc^ in cell culture medium for 24 hours.

### PK digestion of cell lysates

100 µg of total protein, as determined by the BCA assay, was exposed to 10 µg/mL of PK in a final volume of 250 µL at 37 °C for 1 h with shaking at 750 rpm. After digestion, 30 µL of 10x Protease Inhibitor in RIPA buffer was added and the samples were centrifuged for 2 h at 21,147x g at 4 °C. The supernatant was removed, and pellets were resuspended in 30 µL of 1x sample buffer with BME.

### Western blotting

Immunoblotting was performed as previously described (Gatdula et al., 2026). Cells were lysed using RIPA buffer containing 0.1% SDS. Protein normalization was performed using Pierce BCA Protein Assay Kit (Thermo, #23225). Samples are adjusted with 4x Bio-Rad Laemmeli Sample Buffer (2.5% (v/v) β-mercaptoethanol (BME)). Samples are boiled at 100 °C for 10 min and run on a 12% Criterion™ TGX™ Precast Midi Protein Gel for 45 min at 200V. Gels are transferred to a PVDF membrane for 45 min at 200 V using 1x Transfer buffer. Membranes are washed with 0.1 % (v/v) TBS-T for 10 min and blocked for 1 h at RT using a 5% (w/v) Blotto non-fat dry milk dissolved in 0.1% TBST. The membrane is then incubated with primary antibodies diluted in 5% milk with 0.1% TBST overnight at 4 °C. The following day, membranes were washed three times with 0.1% TBST for 5 min each. Membranes were then incubated with a horseradish peroxidase (HRP)-conjugated secondary antibody for 1 h at RT. The membranes were washed again three times before being developed. Quantification of bands were done in ImageJ. The primary antibodies used were human anti-PrP antibody D18 (0.1 µg/ml) and mouse anti-actin antibody (1:10000; Millipore Sigma, A2228). The secondary antibodies used were goat anti-human IgG (H+L) HRP conjugate (1:10000; BioRad, 1721050) and goat anti-mouse IgG (H+L) HRP conjugate (1:10000; BioRad, 1706516).

### PIPLC treatment of cells

Media was removed from cells plated on coverslips and replaced with media containing 2.5 U/mL of PIPLC (Thermo, P6466). Cells were then returned to the incubator and treated for at least 4 hours at 37 °C. Following incubation, PIPLC-media was removed, and cells were washed once with 1X PBS for 5 minutes. Cells are then fixed with 4% PFA in PBS for 12 minutes and denatured with 3M Guanidine HCl in water for 10 minutes. Cells are then washed 5 times with 1X PBS for 5 minutes each. Cells are then processed normally for immunofluorescence and microscopy.

### Immunofluorescence staining and microscopy

Immunofluorescence was performed as described previously (Gatdula et al., 2026). Cells were washed with 1X PBS for 5 min each, followed by fixation in 4% paraformaldehyde (PFA) in PBS for 30 minutes at room temperature. After fixation, cells were washed with PBS, incubated for 5 min with 0.1M glycine in PBS, rinsed, and permeabilized with 0.1% Triton X-100 in PBS for 5 min. The cells were incubated with a blocking buffer (1% w/v BSA) for 30 min, and then probed with primary antibodies for 1 hr. Cells were then washed and incubated with secondary antibodies conjugated to a fluorophore for 1 hr. Cells are then washed three times with PBS for 5 min each and with water for 5 min. Coverslips were then mounted with 5 µl of VECTASHIELD Mounting Medium with DAPI (#H-1000-10) on glass slides. Images were acquired using a Zeiss LSM 700 Laser Scanning Confocal Microscope with 63X oil objectives. The primary antibody used was humanized anti-PrP antibody D18 (10 µg/ml). The secondary antibody was goat anti-human IgG-Alexa Fluor Plus 488 (Invitrogen, cat. # A48276, 1:200).

### Live cell imaging

Live cell imaging was performed using an EVOS FL Auto microscope. Images were collected at 20x magnification and analyzed with the Image J processing package (Fiji).

### MTT assay

After recombinant PrP treatments, media was removed from cells, which were then incubated in the presence of 0.5 mg/mL of MTT in Phenol-red free Opti-MEM (Fischer, #11058021) at 37 °C for at least 2 hours. The solution was removed, and cells were incubated with DMSO at 37 °C for 10 min to dissolve formazan crystals. Absorbance was then read at 540 nm using a BioTek Synergy H1 plate reader. Curves were fit by least-squares regression using GraphPad Prism.

### Purification of recombinant bvPrP (Fig. 6)

Recombinant WT and G127V bvPrPs (residues 23 to 230; Isoleucine at position 109) were expressed and purified as described (Orrú et al., 2015). Briefly, PrP DNA sequences encoding bank vole WT or G127V PrPs were ligated into the Pet41 vector. These were transformed into BL21 *Escherichia coli* (E. coli) and were grown in LB medium in the presence of ampicillin. Protein expression was induced using the autoinduction system (Fox and Blommel, 2009) and was purified from inclusion bodies under denaturing conditions using Ni-NTA superflow resin (Qiagen) with an AKTA fast protein liquid chromatographer. This is followed by refolding with guanidine HCl and eluted using an Imidazole gradient. The eluent was dialyzed into 10 mM sodium phosphate buffer (pH 5.8), filtered in 0.45 µm syringe filter and stored at -80 °C. Concentrations of recombinant PrPs were determined by absorbance at 280 nm using an extinction coefficient of 62,005 M^-1^cm^-1^.

### Recombinant bvPrP treatment of cell lines (Fig. 6D-F)

Medium on CAD5 cells was removed and replaced with medium containing 1.10, 0.55, 0.27, 0.14, or 0 µM of recombinant WT or G127V bvPrPs in Complete Opti-MEM media. The cells were treated for 3 days in an incubator at 37 °C prior to analysis.

### Real-time quaking-induced conversion (RT-QuIC) Assay (Fig. 6C)

RT-QuIC assays were performed as described (Mercer et al., 2025). The reaction mixture contained 10 mM phosphate buffer (pH 7.4), 300 mM NaCl, 0.001% SDS, 1 mM EDTA, 10 μM ThT, and 0.1 mg/mL recombinant WT bvPrP in a final volume of 98 μL in each well of a black, clear bottom, 96-well plate. Reactions were seeded with 2 μL of a 10^-3^ dilution of 22L-infected or uninfected brain homogenate. Plates are incubated in a BMG Polarstar plate reader at 42 °C with a cycle of 1 min shaking at 700 rpm and 1 min of rest. ThT fluorescence measurements were taken every 15 min.

### Graphics

Western blot images were labeled using Lab Figures (formerly Sciugo). Illustrations were made using Bio Render. Statistical bar graphs were made using GraphPad Prism.

## RESULTS

### G126V PrP inhibits prion infection of cultured cells

We first tested whether G126V PrP, the murine equivalent of the kuru-protective polymorphism, could act as a dominant negative inhibitor of prion infection when it is constitutively over-expressed in CAD5 cells. Parental CAD5 cells expressing endogenous levels of WT PrP were stably transfected to co-express G126V PrP under control of a constitutively active CMV promoter. As a control for increased total PrP expression, cells were also transfected with a construct encoding WT PrP. All cells were then exposed to PrP^Sc^ for 24 hours, and the development of chronic infection was assessed by western blotting lysates collected after each passage (6 passages).

Prior to prion infection, transfected CAD5 cells showed stable overexpression and proper cell surface localization of both WT PrP and G126V PrP, which could be released from cells by treatment with phosphatidylinositol-specific phospholipase C (PIPLC) (Fig. 1A-B). As expected, transfected cell lines exhibited a higher amount of total PrP^C^ relative to their untransfected counterpart, which expressed endogenous levels of PrP^C^ (Fig. 1A). Consistent with previous literature, non-transfected CAD5 cells were able to propagate RML and 22L, two mouse-adapted prion strains, based on western blotting for proteinase K (PK)-resistant PrP^Sc^ (Fig 1C). CAD5 cells transfected with the WT PrP construct (CAD5.WT) exhibited the same prion susceptibility, albeit producing higher levels of PrP^Sc^, attributed to the overexpression of WT PrP (Fig. 1D). In contrast, CAD5 cells transfected with the G126V PrP construct (CAD5.G126V) showed reduced amounts of PrP^Sc^ after infection with both RML and 22L (Fig. 1D, E). We conclude from these data that co-expression of G126V PrP with WT PrP significantly impairs establishment of chronic prion propagation in CAD5 cells.

**Fig. 1:**
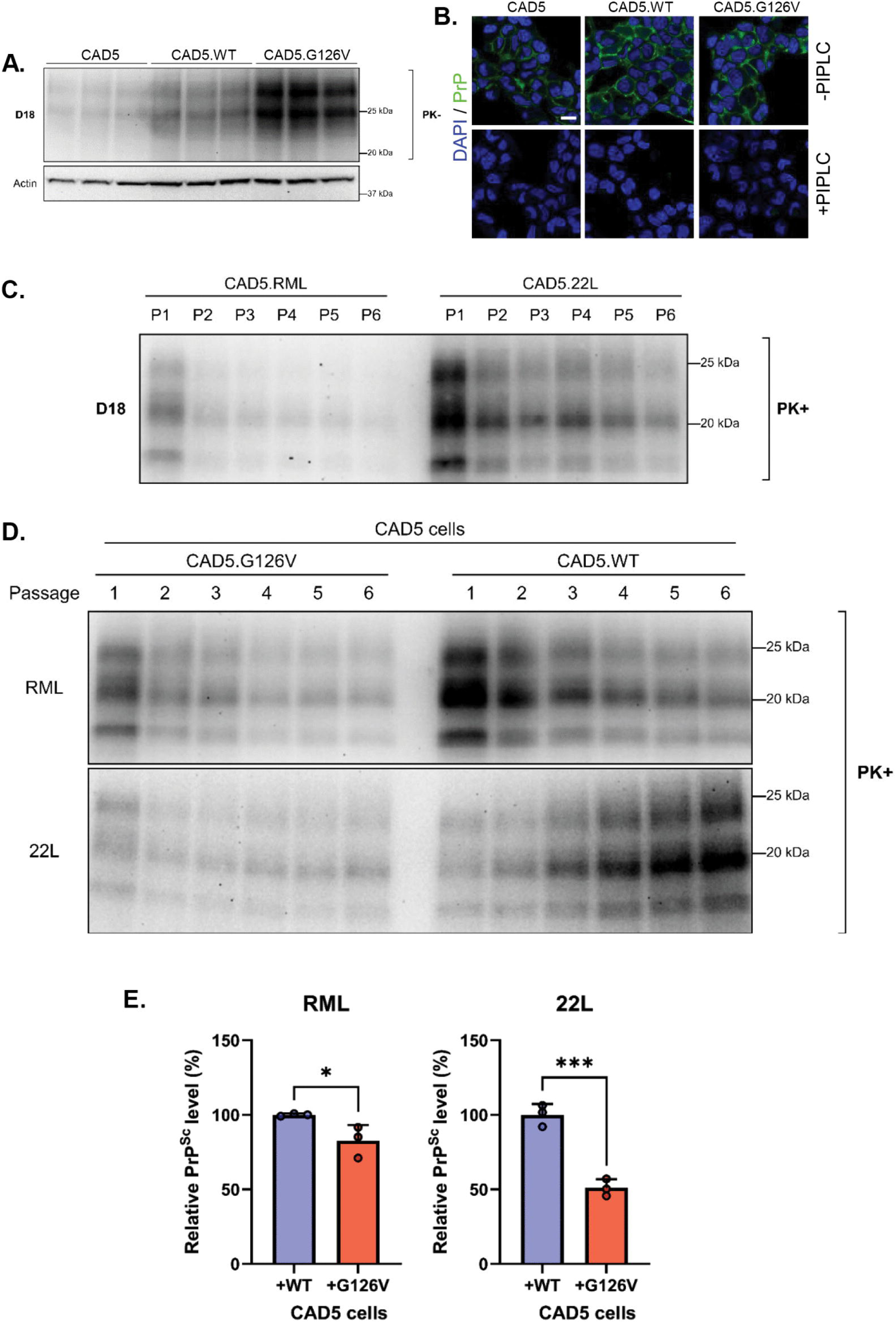
G126V PrP inhibits prion infection of cultured cells. **(A)** Stable expression of WT or G126V PrP^C^ in uninfected CAD5 cells revealed by western blotting with D18 antibody. β-actin was used as a loading control. **(B)** Cell surface localization of PrP in the same cell lines shown in panel A. Uninfected CAD5, either untransfected or expressing WT or G126V PrP, were stained with D18 antibody (green) to detect PrP and with 4′,6-Diamidino-2-phenylindole (DAPI) (blue) to show nuclei. PrP^C^ was released from the cell surface by treatment with PIPLC. Scale bar = 20 µm. Uninfected CAD5 cells, either **(C)** untransfected or **(D)** over-expressing WT or G126V PrP, were exposed to 0.1% RML or 22L brain homogenate for 24h and passaged 6 times to establish a chronic infection. After each passage, cells were lysed, PK digested and immunoblotted for the presence of protease-resistant PrP^Sc^. **(E)** Quantitation of PrP^Sc^ levels of late-stage passages (P5-P6) in Panel D. ImageJ was used to quantify western blots and statistical comparisons were made using a two-tailed t-test. p <0.05 = *; p<0.001 = ***. Bars show the normalized PrP^Sc^ signal relative to CAD5. WT cells set to 100%.

### G126V PrP clears PrP^Sc^ from chronically infected cells

We next tested the ability of G126V PrP to suppress prion infection in chronically infected cells. Constructs encoding WT or G126V PrP were transfected into chronically infected CAD5 cells, selected, and passaged 3 times. Before transfection, prion infection was confirmed by western blotting and immunofluorescence staining, which revealed the presence of PK- and PIPLC-resistant PrP^Sc^ (Fig. 2A-B). As expected, over-expressing WT PrP in chronically infected cells did not alter their ability to propagate prions (Fig. 2C-H). In contrast, over-expressing G126V PrP dramatically decreased the amount of PrP^Sc^ across all three strains (Fig. 2C-H). Expression of G126V PrP caused a large reduction of PrP^Sc^ level in cells infected with 22L, and virtually complete elimination of PrP^Sc^ in cells infected with the RML and ME7 strains. From these results, we conclude that G126V PrP is not only capable of preventing prion infection but is even more effective in eliminating infection in chronically infected cells.

**Fig. 2:**
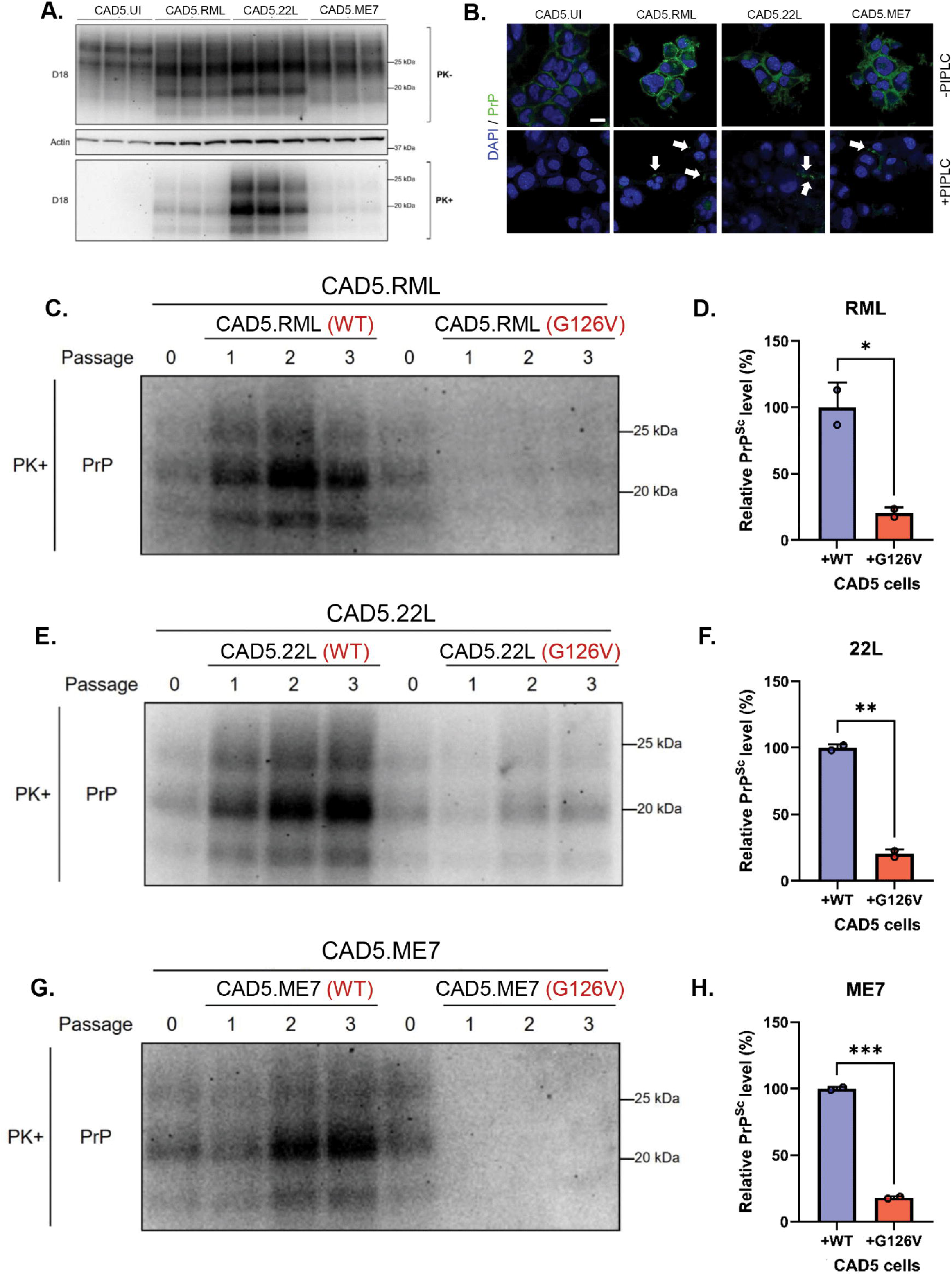
G126V PrP clears PrP^Sc^ from chronically infected cells. **(A)** PrP levels in parental CAD5 cells, either uninfected (UI) or chronically infected with RML, 22L, or ME7 prions revealed by western blotting with D18 antibody. *Upper panels:* Total PrP levels in cell lysates not treated with PK. β-actin was used as a loading control. *Lower panel:* PK digestion of the same cell lysates shown in the upper panels to detect PrP^Sc^. **(B)** Cell surface localization of PrP on the same cells shown in panel A. Chronically infected cells were stained with D18 antibody (green) after PIPLC treatment and GdnHCl denaturation to detect PrP^Sc^ aggregates, and with 4′,6-Diamidino-2-phenylindole (DAPI) (blue) to show nuclei. White arrows point to PrP^Sc^ puncta. Scale bar = 20 µm. **(C, E, G)** CAD5 cells chronically infected with RML, 22L or ME7 prions were transfected with plasmids encoding either WT or G126V PrP, selected under puromycin selection, and passaged 3 times against RML, 22L, and ME7. After each passage, cells were lysed, PK digested and immunoblotted for the presence of protease-resistant PrP^Sc^. Passage 0 shows PK-resistant PrP^Sc^ in chronically infected cell lines before transfection. **(D, F, H)** Quantitation of PrP^Sc^ levels of late-stage passages (P2-3) in Panels C, E, G, respectively. ImageJ was used to quantify western blots, and statistical comparisons were made using a two-tailed t-test. p <0.05 = *; p <0.01 = **; p<0.001 = ***. Bars show the normalized PrP^Sc^ signal relative to CAD5.WT cells set to 100%.

### A doxycycline-inducible system to regulate expression of G126V PrP

Previous observations have shown that hemizygous expression of G127V in transgenic mice prevents infection with some but not all human CJD strains, but that homozygous expression confers complete protection against all tested CJD strains (Asante et al., 2015). This observation suggests that the protective effect depends on the ratio of variant to WT PrP.

To systematically explore this phenomenon in CAD5 cells, we desired a system that would allow us to control and measure expression of G126V PrP independently of endogenous WT PrP, since antibodies are not available that can distinguish the two forms. We therefore designed a doxycycline (Dox)-inducible, bicistronic, lentiviral vector that simultaneously expresses both G126V PrP and eGFP from a tight Tetracycline-Responsive element (tTRE) promoter, using an internal ribosome entry site (IRES2) to allow independent translation of eGFP (Fig. S1A). This system makes it possible to regulate the magnitude and timing of G126V PrP expression using Dox and simultaneously quantitate levels of the variant protein based on eGFP fluorescence. We tested the tunability of the system by generating stably transduced CAD5 cells. Expression of eGFP and G126V PrP was monitored using a fluorescence plate reader, live cell imaging, and western blotting (Fig. S1B-D). We confirmed that CAD5 cells transduced with the construct exhibited a linear correlation between G126V expression and eGFP fluorescence with an R^2^ of 0.946 (Fig. S1E). A major advantage of this system is that it bypasses the requirement for epitope tags to distinguish two PrP proteins expressed in the same cell.

Since Dox has been reported to have an anti-prion effect *in vivo* and *in vitro* (Tagliavini et al., 2000, Cosentino et al., 2005, Forloni et al., 2002, De Luigi et al., 2008, Haïk et al., 2014, Schmitz et al., 2016), we limited the working concentration of Dox in these experiments to ≤ 2 µg/ml, and restricted our analysis to 22L, a prion strain that we have found to be the most resistant to the anti-prion effects of Dox (Fig. S2A-B).

### G126V PrP inhibits initial prion infection in a dose-dependent manner

We then used the Dox-inducible system to test whether G126V PrP could dose-dependently prevent initial prion infection. CAD5 cells expressing the bicistronic construct were treated with either 1 or 2 µg/ml of Dox prior to exposure of the cells to 22L prions; control cells were not treated with Dox (0 µg/ml) (Fig 3A-B). Cell lysates were collected after each of 6 passages following infection and analyzed by western blotting. As expected, control cells not expressing G126V PrP (0 µg/ml Dox) display unimpeded prion propagation, with increasing amounts of PrP^Sc^ after each passage (Fig. 3C). In contrast, cells in which G126V PrP expression was induced with 1 or 2 µg/ml of Dox were restricted in their ability to accumulate PrP^Sc^ during repeated passaging, with this effect being more pronounced in cells induced with 2 µg/ml compared to 1 µg/ml of Dox (Fig. 3C, G). These results demonstrate that the ability of G126V PrP to prevent initial prion infection in CAD5 cells is dose-dependent.

**Fig. 3:**
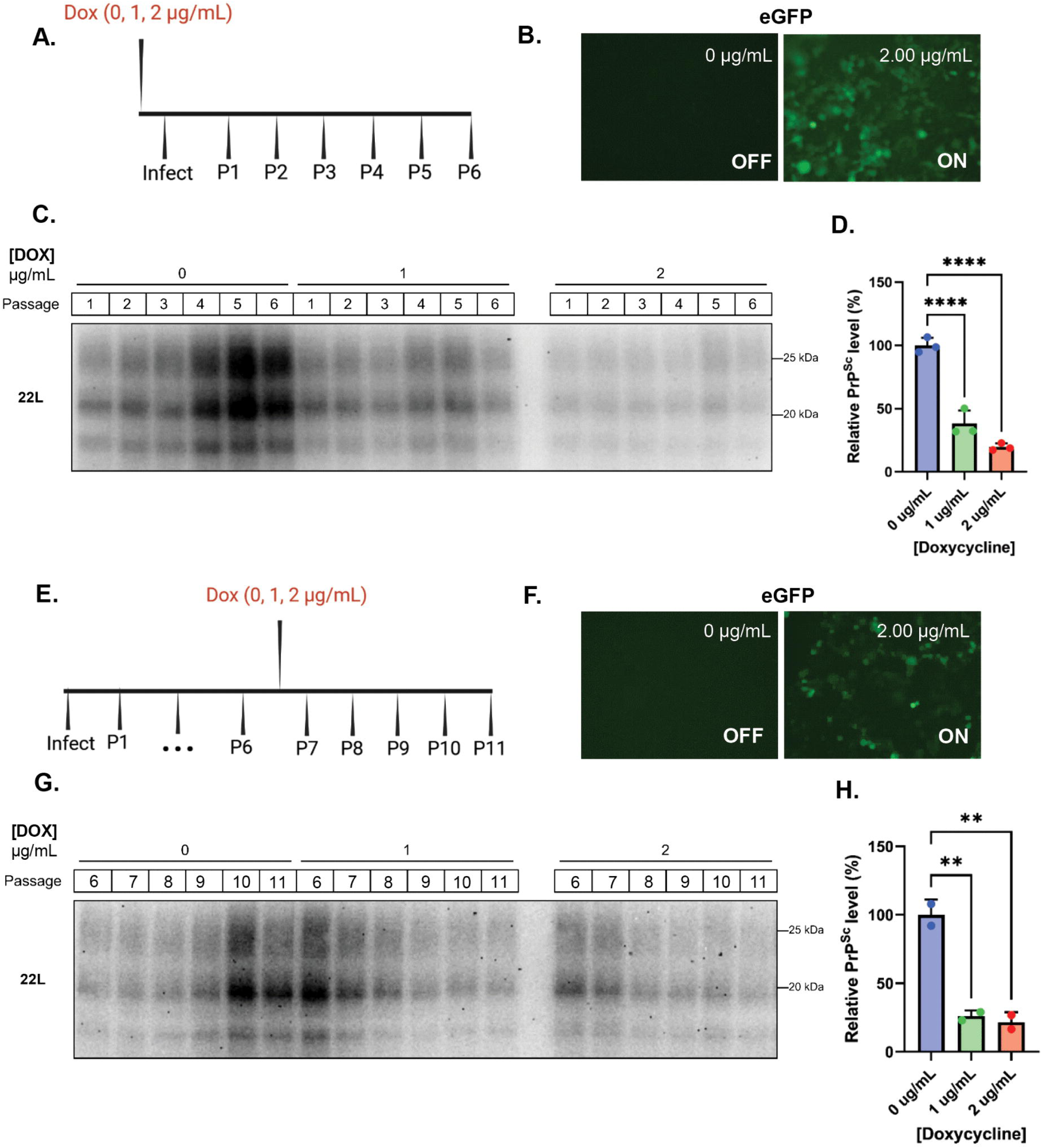
The protective effect of G126V PrP is dose-dependent. **(A)** Timeline of the experiment to demonstrate that G126V PrP inhibits initial prion infection in a dose-dependent manner. Uninfected CAD5 cells that were transduced with the Dox-inducible G126V-IRES2-eGFP construct were induced with 0, 1, or 2 µg/mL of Dox for 3 days before prion infection. Created in BioRender. Gatdula, J. (2026) https://BioRender.com/zx0pt5f **(B)** After 3 days of Dox treatment (2 µg/mL), the same uninfected CAD5 cells were visualized using live-cell imaging to confirm the ON state of G126V expression through eGFP fluorescence. Cells kept at 0 µg/mL of Dox were used as an OFF state negative control. **(C)** After 3 days, induced (1 or 2 µg/mL of Dox) or uninduced (0 µg/mL of Dox) CAD5 cells were exposed to 0.1% 22L brain homogenate for 24h and then passaged 6 times (P1-P6) with our without Dox to establish a chronic infection. After each passage, cells were lysed, PK digested and immunoblotted for the presence of protease-resistant PrP^Sc^. **(D)** Quantitation of PrP^Sc^ levels at passage 6 from western blots like those shown in panel C. ImageJ was used to quantify western blots, and statistical comparisons were made using a one-way ANOVA multiple comparison test. Bars show the normalized PrP^Sc^ signal relative to cells treated with 0 µg/mL of doxycycline set to 100%. Duplicate or triplicate samples were collected from three separate dishes of cells. p<0.0001 = ****; p <0.01 = **. **(E)** Timeline of the experiment to demonstrate that G126V PrP reduces chronic prion infection in a dose-dependent manner. Uninduced (0 µg/mL of Dox) CAD5 cells that were infected in the experiment in Panel A were expanded after passage 6 (P6) and induced with 0, 1, or 2 µg/mL of Dox for 5 additional days (P7-P11). Created in BioRender. Gatdula, J. (2026) https://BioRender.com/zx0pt5f**. (F)** After 3 days of Dox treatment (2 µg/mL), the same CAD5 cells were visualized using live-cell imaging to confirm the ON state of G126V expression through eGFP fluorescence. Cells kept at 0 µg/mL of Dox were used as an OFF state negative control. **(G)** Cells treated with different concentrations of Dox (0, 1, or 2 2 µg/mL) were harvested after passages P6-P11, lysed, PK digested, and immunoblotted for the presence of protease-resistant PrP^Sc^. Passage 6 represents PrP^Sc^ in chronically infected cell lines before induction. **(H)** Quantitation of PrP^Sc^ levels at passage 11 from western blots like those in panel G. Quantitation was performed as described in panel D.

### G126V PrP reduces chronic prion infection in a dose-dependent manner

We then used the Dox system to test whether the curative effect of G126V on chronically infected cells is also dose dependent. Cells expressing the bicistronic construct were first chronically infected with 22L for six passages in the absence of Dox and were then exposed to either 1 or 2 µg/ml of Dox for six more passages; control cells were not treated with Dox (0 µg/ml) (Fig. 3D-E). Cell lysates were collected after each of the six passages following Dox induction and analyzed by western blotting. Paralleling its ability to inhibit initial infection, G126V PrP was also able to suppress chronic infection, and this effect was also dose-dependent (Fig. 3F, H).

### G127V PrP protects against a broad range of prion strains

Our experiments thus far were restricted to CAD5 cells expressing murine G126V PrP that were infected with mouse-adapted prion strains. To test whether the inhibitory effect of the variant PrP is broadly applicable to other prion strains, we used cells expressing bank vole PrP (bvPrP), which is a “universal substrate” that can be converted to PrP^Sc^ by a wide range of biologically derived and synthetic prion strains from multiple species (Watts et al., 2014, Orrú et al., 2015, Mok et al., 2021). A plasmid encoding WT bvPrP was first transfected into CAD5 cells in which the endogenous PRNP gene had been knocked-out by CRISPR-Cas9 editing. After selection, these cells were then transfected with plasmids encoding either WT bvPrP or G127V bvPrP (bank vole numbering) carrying a different selection marker. Following a second selection, these cells were then exposed to naturally derived prion strains from different species (mouse, hamster, elk, and deer) and to synthetic, recombinant prion strains (mouse) for 24 hours, and chronic infection was assessed by western blotting after passaging 8-9 times.

Prior to infection, transfected CAD5 cells showed stable overexpression and proper cell surface localization of both WT bvPrP and G127V bvPrP (Fig. 4A-B). Similar to previous experiments (Fig. 1A), an increased level of PrP^C^ expression was observed in the co-expression cell lines (Fig. 4B). We first tested the susceptibility of these lines to the 22L strain. Consistent with previous literature (Arshad et al., 2023), CAD5.*Prnp^-/-^* cells reconstituted with WT bvPrP were able to propagate the 22L strain while the G127V mutant was completely refractory to propagation of 22L prions (Fig. 4C). WT bvPrP co-expression cell lines (WT+WT) showed an increased amount of PrP^Sc^, but the PrP^Sc^/PrP^C^ ratio was indistinguishable from singly transfected cells expressing WT bvPrP (Figs. 4C and S3A). G127V bvPrP co-expression cell lines (WT+G127V) showed a statistically significant decrease in PrP^Sc^ relative to WT+WT cells (Figs. 4C and S3A).

**Fig. 4:**
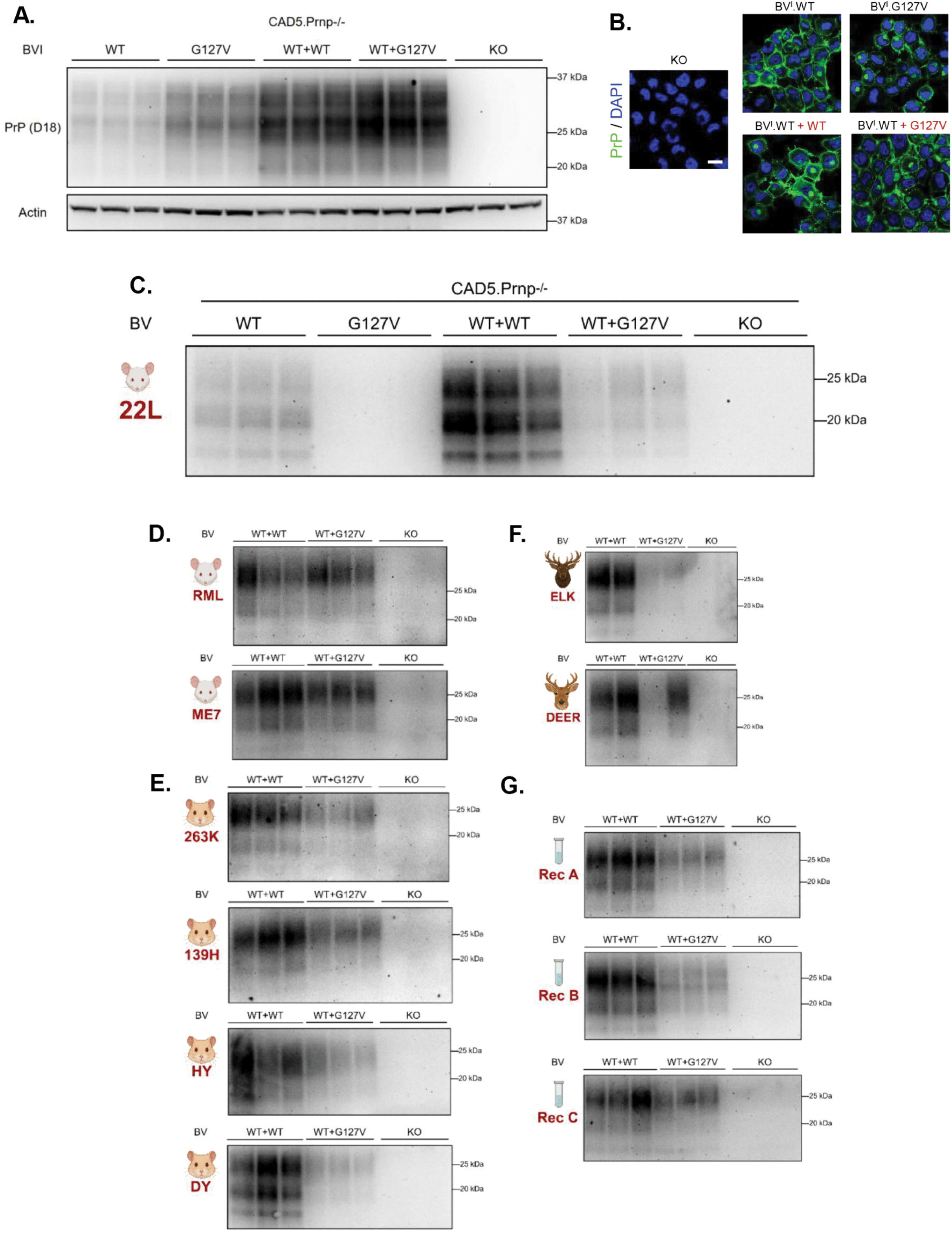
G127V PrP protects against a broad range of prion strains. **(A)** Stable expression of WT or G127V bvPrP^C^ in CAD 5 cells. Uninfected CAD5.Prnp^-/-^ cells, either untransfected (KO), expressing only WT or G127V PrP, or co-expressing both WT and G127V PrP were analyzed by western blotting with D18 antibody. β-actin was used as a loading control. **(B)** Cell surface localization of WT or G127V PrP^C^ expressed in the same CAD5 cell lines shown in panel A. Cells were stained with D18 antibody (green) to detect PrP and with 4′,6-Diamidino-2-phenylindole (DAPI) (blue) to show nuclei. Scale bar = 20 µm. **(C)** The indicated CAD5 cell lines were exposed to mouse 22L brain homogenate for 24h and passaged 8-9 times to establish a chronic infection and dilute out brain inoculum. After P9, cells were lysed, and the lysates PK digested and immunoblotted to detect PrP^Sc^. **(D-G)** CAD5 cell lines (KO [knock-out], WT+WT and WT+G127V) were exposed for 24 hrs. to the following inocula: brain homogenates from RML or ME7 infected mice **(D)**; brain homogenates from 263K, 139H, HY, or DY infected hamsters **(E)**; cell homogenates from deer or elk cell lines infected with Chronic Wasting Disease (CWD) prions **(F)**; and three preparations of recombinant prions **(G)**. After prion exposure, cells were passaged 8-9 times, lysed, and the lysates PK digested and immunoblotted to detect PrP^Sc^. Duplicate or triplicate lanes represent samples collected from two or three separate dishes of cells. Knockout CAD5 cells were used as negative controls. Species images were created in BioRender. Gatdula, J. (2026) https://BioRender.com/wrcdecz.

We then tested two other mouse-derived strains (ME7 and RML), as well as brain-derived prion strains from other species, and found that co-expression of G127V bvPrP generally reduced the accumulation of PrP^Sc^ (Figs. 4D-G and S3B-E). The reduction ranged from 30-70% for the mouse strains ME7 and 22L (Figs. 4D and S3B); 50-80% for the hamster strains 139H, 263K, DY (Drowsy), and HY (Hyper) (Figs. 4E and S3C); and 80% for an elk prion strain (Figs. 4F and S3D). The mouse RML strain and a deer prion strain showed a decrease in PrP^Sc^ that was not statistically significant (Figs. 4D, 4F, S3B, and S3D).

Recently, it has become possible to generate a wide range of synthetic prion strains from multiple species using recombinant PrP in an *in vitro* amplification system (Eraña et al., 2024, Pérez-Castro et al., 2025). We tested whether co-expression of G127V bvPrP inhibited infection by three such strains. Recombinant A (stML-01-Dx), B (btML-89-Dx), and C (btML-90-Dx) are recombinant mouse prion preparations generated spontaneously by Protein Misfolding Shaking Amplification (PMSA) in the presence of sulfated dextran as described previously (Pérez-Castro et al., 2025) that display strain-specific characteristics when inoculated into mice. We found that these strains are also capable of infecting CAD5 cells expressing bvPrP (Fig. 4G). Paralleling its effect on biologically derived prion strains, G127V bvPrP reduced PrP^Sc^ accumulation by 30-50% when co-expressed with WT bvPrP in CAD5 cells following exposure to each of the three synthetic prion strains (Figs. 4G and S3E). Thus, the G127V PrP variant is able to inhibit prion infection by a wide range of biologically derived and synthetic prion strains.

### Inhibition of prion propagation persists when expression of G126V PrP is turned off

To test the persistence of the G126V PrP inhibitory effect, we utilized the Dox inducible system to shut off expression of G126V PrP in previously infected cells. We anticipated that this would allow residual prions to begin propagating again. CAD5 cells carrying the bicistronic G126V PrP plasmid were induced with 2 µg/mL of Dox starting before initial infection, as in Fig. 3A. The expression of G126V was then turned off by removing Dox from the system (0 µg/mL) (Fig. 5A-B). These cells were passaged seven times and cell lysates were collected from each passage to monitor changes in the amount of PrP^Sc^. Surprisingly, PrP^Sc^ levels did not rebound during passaging in the absence of Dox and did not return to the levels seen in infected cells never treated with Dox (Fig. 5C). Monitoring eGFP fluorescence and the reduction of total PrP levels detected through western blot confirmed that the bicistronic system had been turned off (Fig. 5B and 5D).

**Fig. 5:**
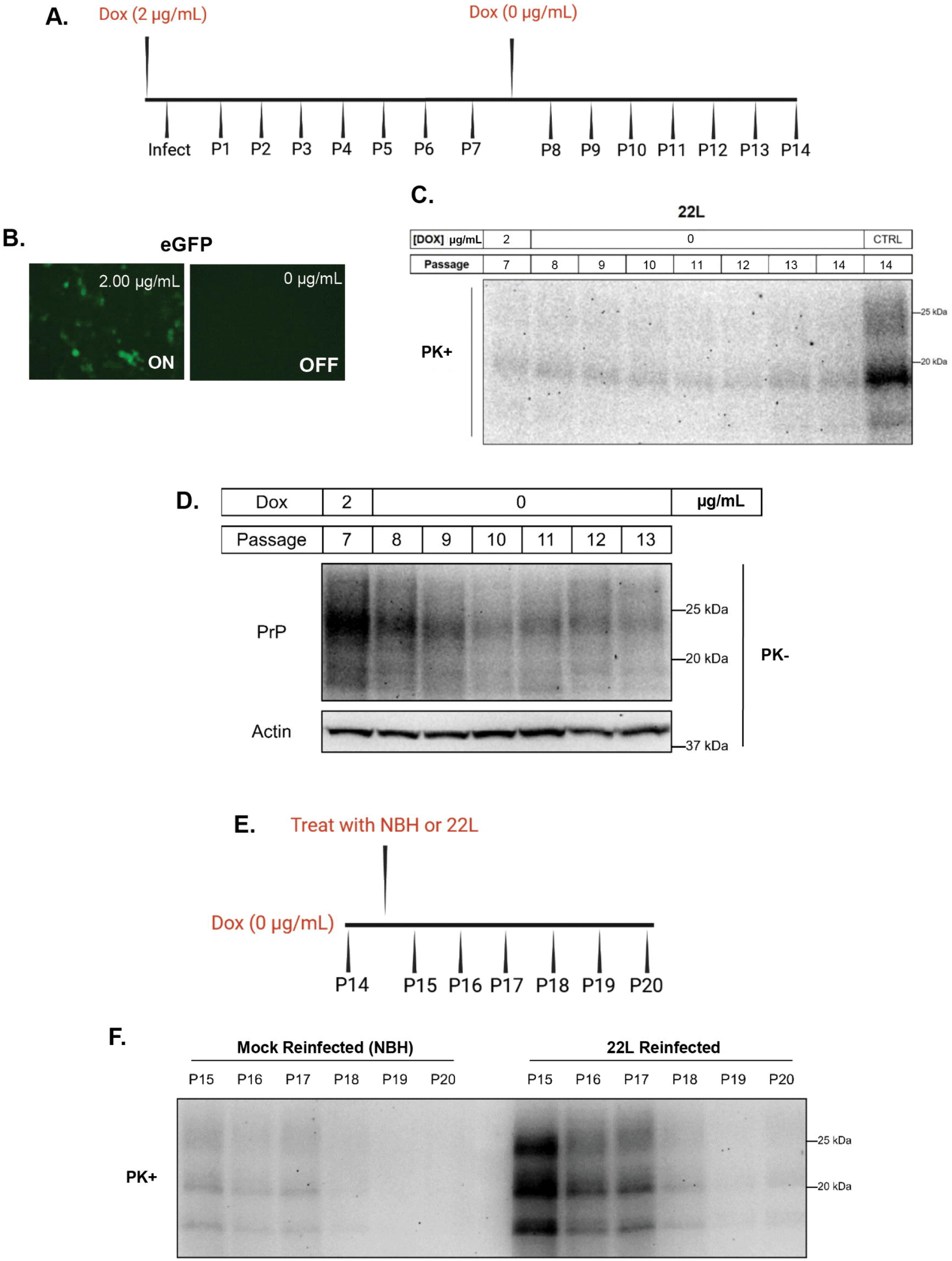
Inhibition of prion propagation persists when expression of G126V PrP is turned off. **(A)** Timeline of the experiment. Cells induced to express G126V PrP by treatment with 2 µg/mL Dox were infected with 22L prions. After 7 passages (P7), the cells were switched to medium without Dox 0 µg/mL to shut off expression of G126V PrP. Cells were then passaged 7 more times (P8-P14). Created in BioRender. Gatdula, J. (2026) https://BioRender.com/zx0pt5f **(B)** Three days after removing Dox at P7, cells were visualized using live-cell imaging to confirm the OFF state (0 µg/mL) of G126V expression through eGFP fluorescence. Cells kept at 2 µg/mL of Dox were used as an ON state positive control. **(C and D)** After each passage (P7-P14), cells were lysed, after which the lysates were either digested with PK **(C)** or were left undigested **(D)**, followed by immunoblotting to reveal PrP^Sc^ or total PrP, respectively. The sample in the right-hand lane of Panel C (labeled *CTRL*) is from control cells that have never been induced with Dox. P7 represents cells induced with Dox from P1-P7 before removal of Dox at P8. **(E)** Timeline of the experiment. After 14 passages, cells switched to Dox-free medium at P7 were exposed to 1% 22L brain homogenate or 1% normal brain homogenate (NBH) for 24 h and then passaged 6 more times (P15-P20) in the absence of Dox to determine if the cells can be reinfected to boost PrP^Sc^ levels. **(F)** After each passage, cells were lysed, PK digested and immunoblotted for the presence of protease-resistant PrP^Sc^.

We then asked whether re-infecting these cells with 22L infected brain homogenate (IBH) or normal brain homogenate (NBH) for 24 hours (Fig. 5E) could enhance levels of PrP^Sc^. Cells were passaged for six more times after re-infection and cell lysates were collected from each passage for western blotting. Surprisingly, CAD5 cells in which PrP^Sc^ were suppressed by expression of G126V PrP did not display increased PrP^Sc^ levels after re-exposure to a high concentration of 22L prions (Fig. 5F).

### Recombinant G127V PrP reduces PrP^Sc^ in chronically infected cells

Thus far, we have demonstrated the ability of mouse G126V PrP or bank vole G127V PrP to act as dominant-negative inhibitors of PrP^Sc^ propagation when co-expressed with WT PrP in cultured cells. We wondered whether a non-membrane-anchored, recombinant form of the variant protein applied extracellularly might have the same effect. If so, this would enhance its ability to spread within the CNS in a clinical setting.

We generated recombinant WT and G127V bvPrPs by expression in *E. coli* and confirmed their purity by SDS-PAGE and western blotting (Fig. 6A-B). We also tested the ability of the recombinant proteins to serve as substrates using the Real Time Quaking Induced Conversion (RT-QuIC) assay seeded with 22L PrP^Sc^ (Fig. 6C). As expected, recombinant G127V PrP showed impaired conversion compared to relative WT PrP. To test whether recombinant G127V exhibits a dominant-negative effect, we treated 22L-infected CAD5 cells expressing WT bvPrP with recombinant G127V PrP for 3 days and then assayed PrP^Sc^ levels by western blotting (Fig. 6D). We found that recombinant G127V PrP reduced PrP^Sc^ levels in a dose-dependent manner, with an IC50 value of 1.276 µM (Fig. 6F). In contrast, recombinant WT bvPrP had no effect on PrP^Sc^ levels (Fig. 6E, F). To check for cytotoxicity of the recombinant proteins, we treated 22L-infected CAD5 cells for 3 days with a range of concentrations and performed a 3-(4, 5-dimethylthiazol-2-yl)-2, 5-diphenyltetrazolium bromide (MTT) assay (Fig. 6F).

**Fig. 6:**
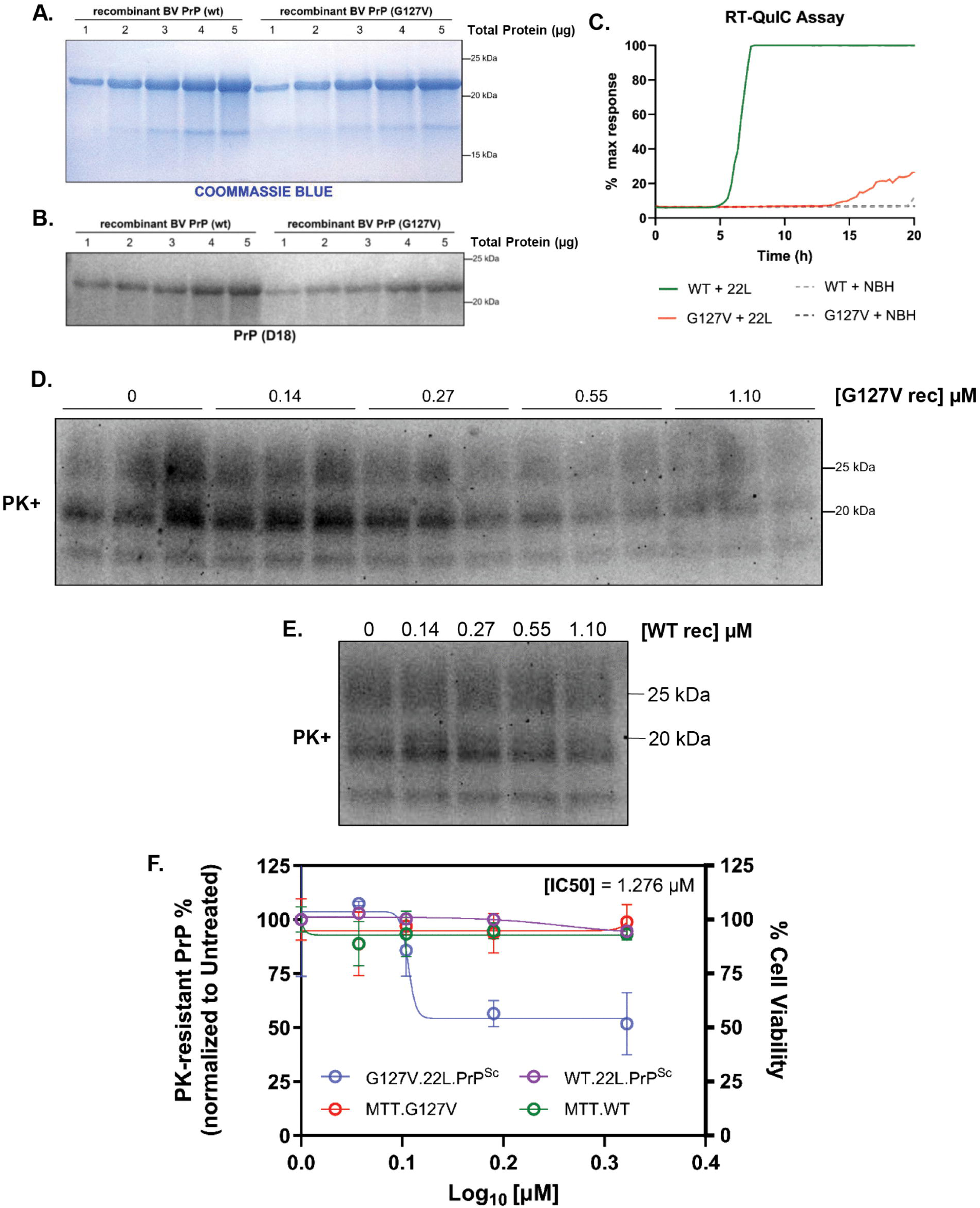
Recombinant G127V PrP reduces PrP^Sc^ in chronically infected cells. **(A)** Coommassie blue stained SDS-PAGE gel staining of indicated amounts of recombinant WT and G127V bvPrP. **(B)** The indicated amounts of recombinant WT and G127V bvPrP were western blotted and probed the D18 antibody. **(C)** RT-QuIC assays testing conversion of 0.1 µg of recombinant WT or G127V bvPrP after seeding with 10^-3^ dilution of of 22L-infected brain homogenate (*22L*) or normal brain homogenate (*NBH*). Each condition was run in quintuplicate. 22L-infected CAD5 cells expressing bvPrP were treated with different concentrations of recombinant G127V **(D)** and WT **(E)** bvPrP (1.10, 0.55, 0.27, 0.14, and 0 µM) for 3 days, followed by immunoblotting of cell lysates to detect protease-resistant PrP^Sc^. Parallel wells were assayed for cytotoxicity using MTT. Each condition was run in triplicate. **(F)** PrP^Sc^ levels and MTT measurements of cell viability were quantitated and expressed as a percentage of the values for untreated cells.

## DISCUSSION

In this study, we use cell-based infection models to explore the use of a naturally occurring, kuru-protective *PRNP* variant (G127V) as an alternative therapeutic strategy to treat prion diseases. We demonstrated that co-expression of PrP harboring the G127V substitution or its murine homologue G126V has a dose-dependent, dominant negative effect on infection and subsequent propagation of a broad range of prion strains, both biologically derived and synthetic. Surprisingly, we discovered that this inhibitory effect was sustained after G126V PrP expression was terminated, raising interesting mechanistic questions with implications for use of this variant in a therapeutic setting. Finally, we report that an anchorless, recombinant version of G127V PrP is capable of reducing PrP^Sc^ propagation when applied to cells externally, suggesting the feasibility of using this form as a biologic therapeutic (Fig. S4).

### Structural effects of the G127V polymorphism

The wide strain and species specificity of the G127V effect on prion infection and propagation we have demonstrated here indicates that this variant PrP engages a highly conserved, inhibitory mechanism in which the PrP molecule is conformationally locked in a PrP^C^ state that impedes its transition to the PrP^Sc^ state. Structural studies and molecular dynamics simulations suggest that this substitution constrains the polypeptide backbone and alters PrP dimerization during the initial events of misfolding (Hosszu et al., 2020, Sangeetham et al., 2021, Zheng et al., 2018, Zhou et al., 2016). The G127V polymorphism has also been shown to impede fibrilization of recombinant PrP (Arshad et al., 2023, Hosszu et al., 2020, Huang et al., 2020, Sabareesan and Udgaonkar, 2017, Lee et al., 2007), a finding which we have confirmed here (Fig. 6C).

### Dominant negative effect of the G127V polymorphism

Previous experiments on transgenic mice co-expressing different ratios of WT and G127V PrP and inoculated with CJD and kuru isolates provided evidence for a dose-dependent, dominant negative inhibitory effect of the G127V variant (Asante et al., 2015). We have demonstrated that the same phenomenon occurs in a cell culture setting using an inducible expression system that permits expression of varying amount of G126V PrP in proportion to WT PrP. We found that prophylactic expression of G126V PrP in CAD5 cells prior to infection substantially reduced prion propagation in a dose-dependent manner. We also demonstrated that G126V PrP dose-dependently suppresses prion propagation in cells with established infection.

These results raise the interesting question of the molecular mechanism underlying the dominant negative effect of the G127V variant. One possibility is that G127V PrP, which is locked in a PrP^C^ conformation, competes with WT PrP for incorporation into PrP^Sc^ fibrils, possibly at their growing ends, thereby reducing the efficiency with which PrP^Sc^ converts new molecules of PrP, or perhaps completely preventing further elongation. Alternatively, PrP^Sc^ molecules that have incorporated the G127V variant may be less stable, and fragment more easily, contributing to their faster clearance from the cell by degradative processes. Single-molecule analyses of PrP^Sc^ fibril growth are consistent with both of these suggestions (Sun et al., 2023, Sang et al., 2018). Several older studies have described a dominant-negative effect on PrP^Sc^ propagation in cells and transgenic mice of PrP molecules carrying positively charged amino acid substitutions at residues 167, 171, or 218 (mouse PrP numbering), which were postulated to form a binding site for a hypothetical co-factor called “protein X” (Perrier et al., 2002, Perrier et al., 2000, Kaneko et al., 1997). Moreover, recombinant mouse PrP containing a Q→K substitution at codon 218, the analogous position as a human E219K polymorphism found in the Japanese population and believed to confer resistance to CJD (Furukawa et al., 1995, Seno et al., 2000, Shibuya et al., 1998), inhibited PrP^Sc^ propagation in N2a cells (Kishida et al., 2004). In principle, all of these effects could be explained by direct competition of the mutant PrP molecules for growth sites on PrP^Sc^ fibril ends or by destabilization of the fibrils, as suggested above for G127V PrP, without having to postulate the involvement of an accessory protein X molecule.

### Persistence of G126V inhibitory effect

Our results have uncovered a surprising property of G126V PrP: its dominant-negative suppression of prion propagation persists even after synthesis of the variant protein is turned off. Using the dox-regulated expression system, we found that removal of dox to turn off G126V synthesis in prion-infected cells did not result in rebound of PrP^Sc^ levels to those seen in the absence of dox, even after multiple passages (Fig. 5). Moreover, re-exposure of the cells without dox to a second round of prion infection did not result in increased levels of PrP^Sc^. In both cases, residual PrP^Sc^ was present at the time of dox removal, and it might be expected that these molecules would have propagated further over ensuing passages in the absence of G126V PrP. It is possible that synthesis of G126V PrP was not completely shut off by dox removal due to leaky transcriptional control, or that small numbers of G126V PrP molecules remained bound to existing PrP^Sc^ fibrils. However, these scenarios seem unlikely since sub stoichiometric ratios of residual G126V to WT PrP would have minimal inhibitory effect, and residual G126V molecules bound to PrP^Sc^ fibrils would have been diluted out with continual passage.

Instead, we suggest two alternative explanations for the persistence of the G126V inhibitory effect. First, the conversion-incompetent G126V molecules might have caused a persistent change in the structure of the PrP^Sc^ fibrils into which they were incorporated, resulting in a more slowly propagating strain. This process would be analogous to the well-known phenomenon of prion strain adaptation (Block and Bartz, 2023). Alternatively, the presence of G126V PrP might have induced long-lasting cellular changes that result in slower prion replication or enhanced degradation, for example shunting of PrP to intracellular compartments less favorable for conversion or increased activity of lysosomes and/or autophagosomes. Further work will be necessary to distinguish between these possibilities.

### G127V PrP as a therapeutic

At present, therapeutic efforts to treat prion diseases have largely focused on three strategies: (1) inhibiting PrP^C^ conversion to PrP^Sc^ using small molecules, antibodies, or other ligands that bind to PrP^C^ (Yamaguchi et al., 2019, Nicoll et al., 2010, Masone et al., 2023, White et al., 2003, Mead et al., 2022, Mercer et al., 2024, Mercer and Harris, 2019, Trevitt and Collinge, 2006); (2) reducing the synthesis of PrP^C^, using genetic tools (Chou et al., 2025, Minikel et al., 2020); and (3) accelerating the clearance of PrP^Sc^ by manipulation of cellular degradative pathways (Mercer et al., 2025). Of these strategies, reduction PrP^C^ has progressed most rapidly to clinical application, in part because of the undisputed role of PrP^C^ in prion propagation, as well as recent success in extending survival in preclinical models of prion disease. However, complete ablation of PrP^C^ expression in PrP-knockout mice has been associated with subtle developmental and behavioral abnormalities and peripheral neuropathy (Coitinho et al., 2003, Collinge et al., 1994, Carleton et al., 2001, Prestori et al., 2008, Beraldo et al., 2011, Criado et al., 2005, Nishida et al., 1997), raising concerns about the long-term risks of PrP^C^ reduction in humans, and emphasizing the need for additional therapeutic approaches.

Our findings establish a mechanistic and cell biological framework for developing the naturally protective G127V PrP polymorphism as a novel therapeutic reagent for treatment of human prion diseases. G127V PrP is an effective inhibitor of prion infection and propagation when expressed in a cellular context along with WT PrP, or when administered extracellularly as a recombinant protein. Therefore, in terms of potential modes of delivery, G127V PrP would offer several options, including administration of the soluble protein as a biologic or via production in genetically engineered cells, or using viral vectors programmed to synthesize the membrane-anchored or anchorless version. It remains to be determined whether the protective effect of G127V PrP extends to PrP molecules carrying mutations linked to familial prion diseases. It will also be worthwhile to explore whether other structurally constraining variants of PrP would offer even better protection than G127V, although a considerable advantage of the latter is that it is a naturally occurring polymorphism in humans.

## Supporting information

Supplemental Material: Raw Western Blots

## CRediT authorship contribution statement

**Jean R.P. Gatdula:** Writing – review and editing, Writing – original draft, Visualization, Validation, Software, Methodology, Investigation, Formal analysis, Data curation, Conceptualization. **Isabel C. Orbe:** Visualization, Methodology, Data curation. **Samantha G. Tolton:** Methodology. **Linnea M. Saunders:** Methodology. **Janelle S. Vultaggio:** Methodology. **Robert C.C. Mercer:** Methodology, Writing – review and editing, Supervision. **Jason C. Bartz:** Resources, Writing – review and editing. **Glenn C. Telling:** Resources, Writing – review and editing. **Hasier Eraña:** Methodology, Resources, Writing – review and editing. **Joaquín Castilla:** Methodology, Resources, Writing – review and editing. **David A. Harris:** Writing – review and editing, Supervision, Funding acquisition.

## Data availability

All data supporting the findings of this study are available within the article and its supplementary information files. Submission contains all raw data required to replicate the results of the study.

## Supplementary Data

Raw western blot files are included as an extra supplemental file.

## ACKNOWLEDGEMENTS

We thank Mikel Garcia-Marcos (Boston University) for providing Lenti-X HEK293T cells and packaging plasmids (psPAX2 and pMD2.G); Joel Watts (University of Toronto) for wild type CAD5 and CAD5.Prnp^-/-^ cells and pcDNA.3-G418 WT bvPrP (I109); and Byron Caughey for the pET41 plasmid.

## SUPPORTING INFORMATION

**Fig. S1:**
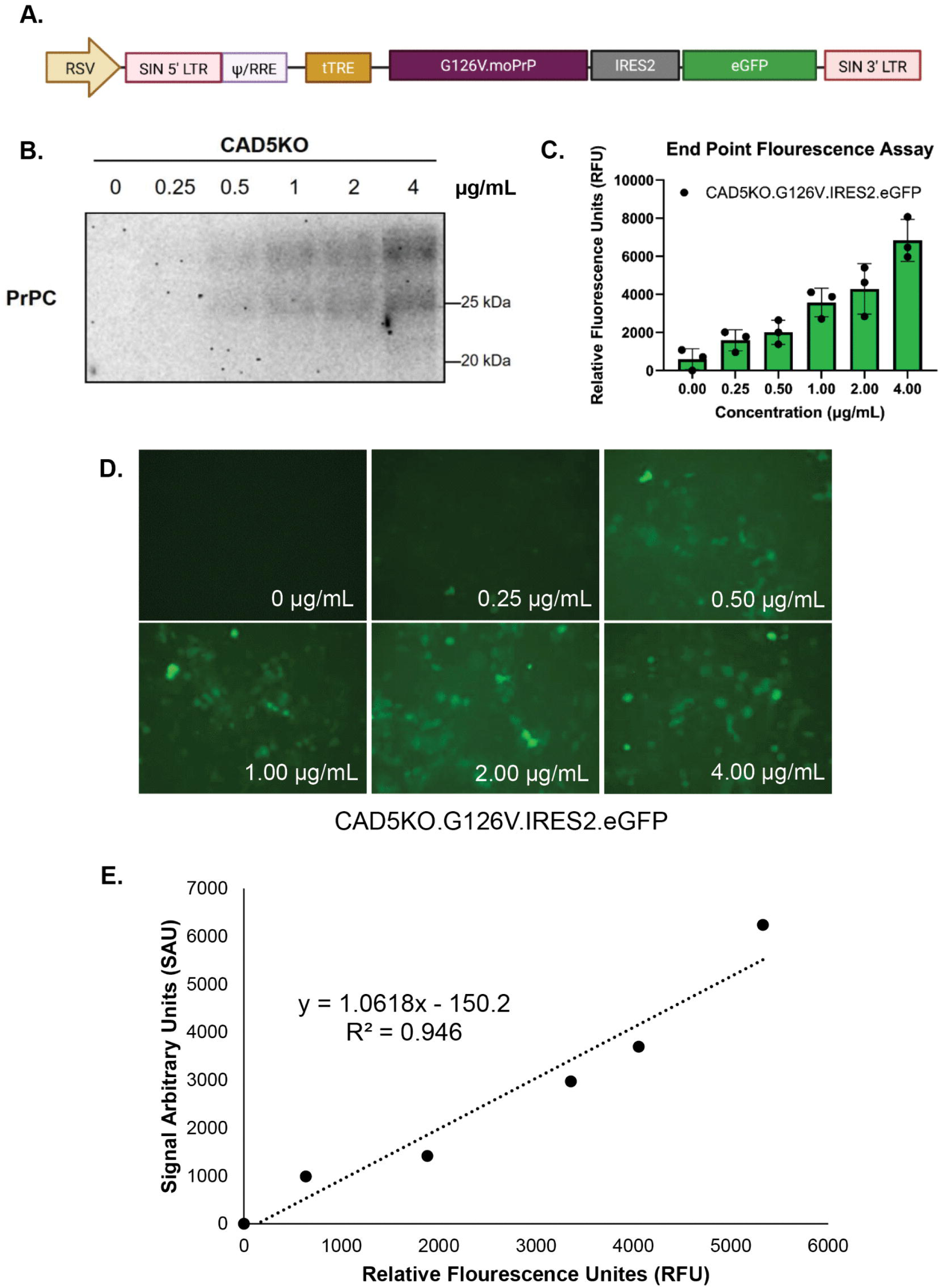
A doxycycline-inducible system to regulate expression of G126V PrP. **(A)** Map of the Dox-inducible, bicistronic, lentiviral vector that simultaneously expresses both G126V mouse PrP and eGFP from a tight Tetracycline-Responsive element (tTRE) promoter, with an internal ribosome entry site (IRES2) to allow independent translation of eGFP. Created in BioRender. Gatdula, J. (2026) https://BioRender.com/sldyc7e **(B)** Uninfected CAD5.Prnp^-/-^ that were stably transduced with the construct were treated with 0, 0.25, 0.5, 1.0, 2.0, and 4.0 µg/mL of Dox for 3 days, lysed, and immunoblotted for PrP using D18. **(C)** The transduced cells were plated in a 96-well format and induced with the same Dox concentrations as in panel B. After 3 days of treatment, eGFP fluorescence was measured using a BioTek Synergy H1 Plate Reader with an excitation wavelength of 485 nm and emission wavelength of 528. **(D)** In parallel, the same cells as in panel C were visualized using a live cell imaging system to confirm the dose-responsiveness of expression from the tTRE promoter. **(E)** The PrP immunoblot signals in Panel B were plotted against the eGFP fluorescence signals in Panel C, demonstrating a linear relationship between G126V and eGFP levels.

**Fig. S2:**
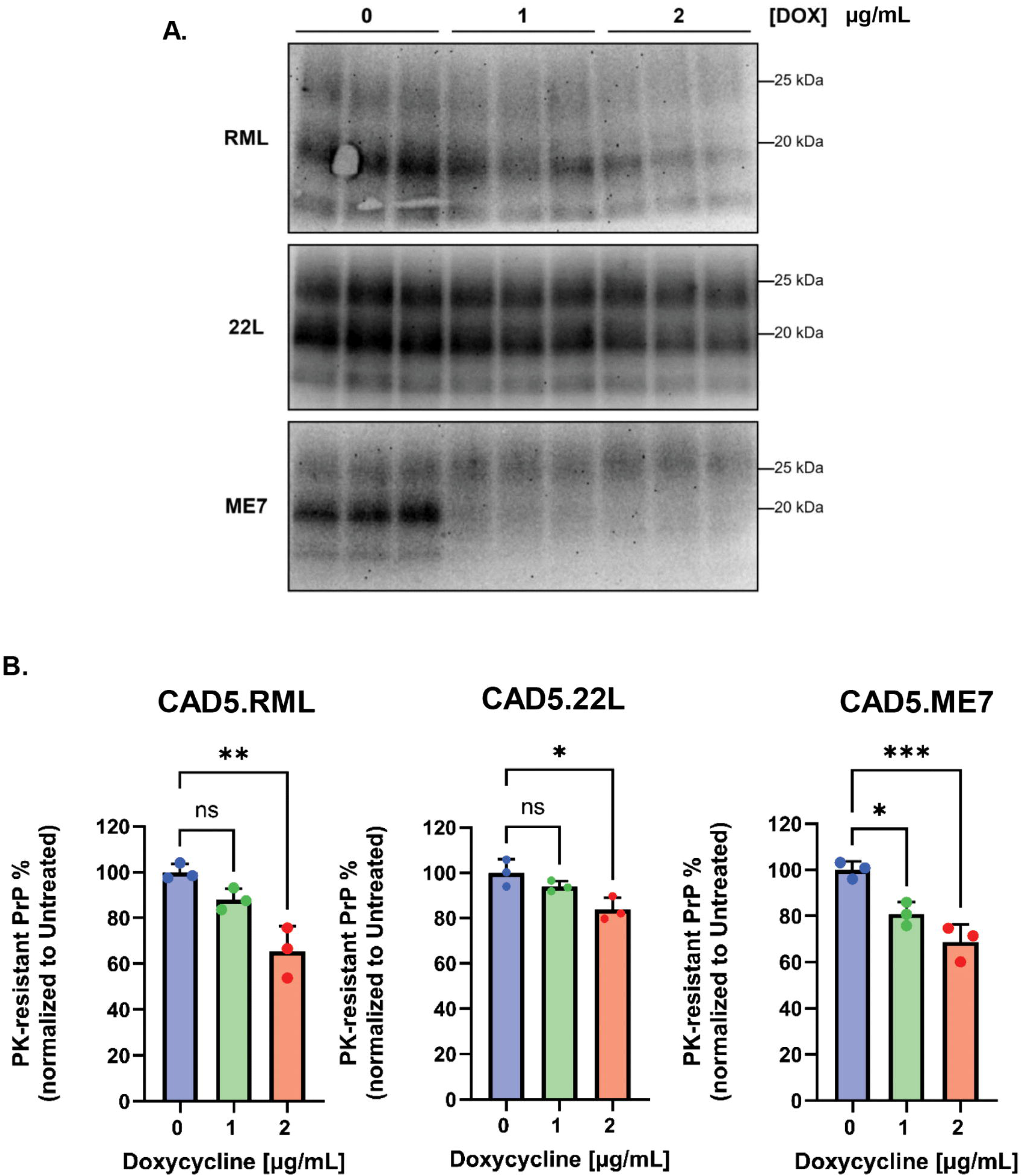
22L strain is relatively resistant to the anti-prion effects of Dox in CAD5 cells. **(A)** CAD5 cells were exposed to 0.1% 22L, RML, and ME7 brain homogenate for 24h and passaged 6 times in the presence of 0, 1, or 2 µg/mL of Dox. After P6, cells were lysed, PK digested and immunoblotted for the presence of PrP^Sc^. Triplicate lanes represent samples collected from three separate dishes of cells. **(B)** Quantitation of PrP^Sc^ levels in Panel A. ImageJ was used to quantify western blots and statistical comparisons were made using a one-way ANOVA multiple comparison test. For all statistical analyses, bars show the normalized PrP^Sc^ signal relative to untreated WT cells set to 100%. p<0.001 = ***; p <0.01 = **; p <0.05 = *; ns = not significant.

**Fig. S3:**
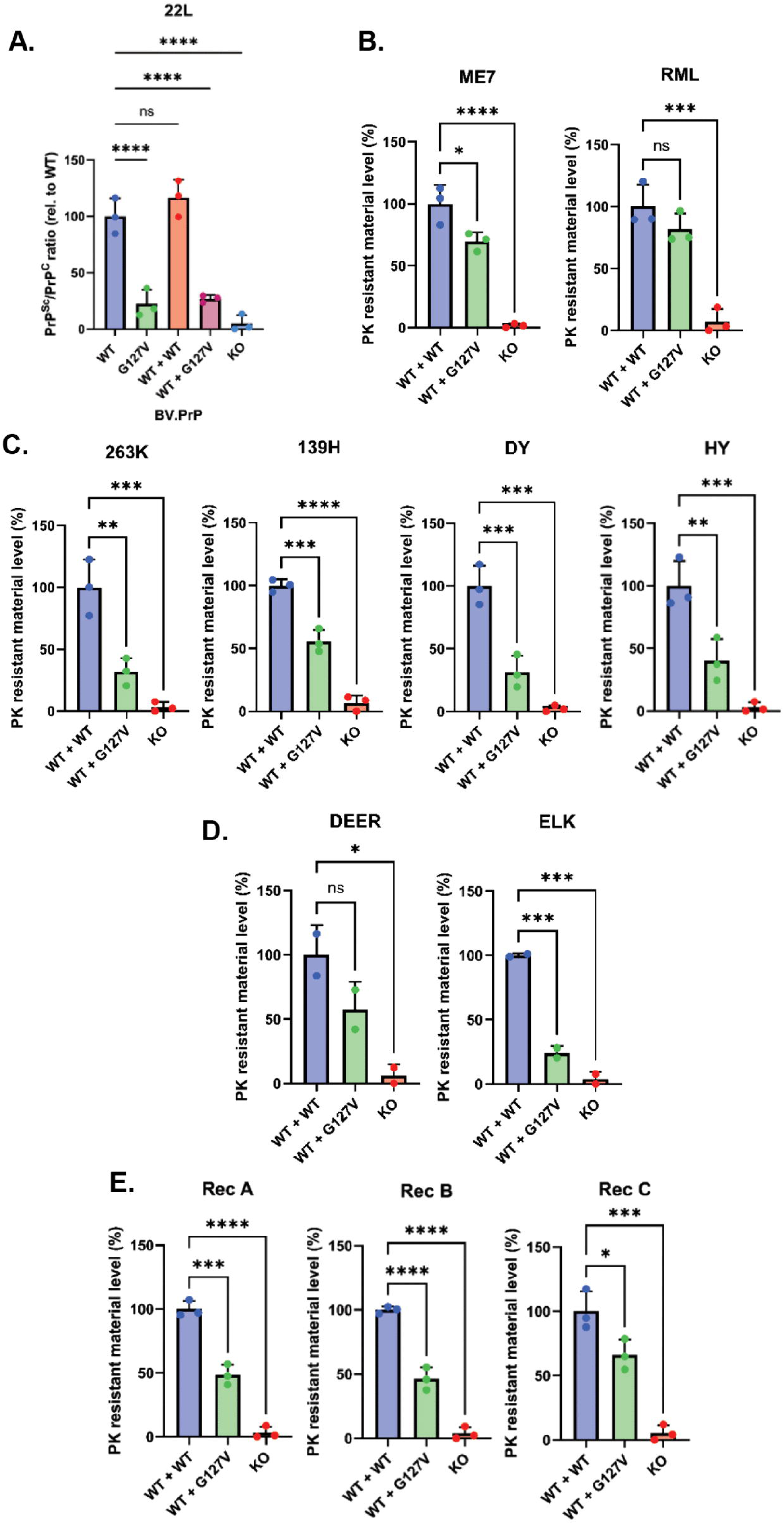
Quantitation of PrP^Sc^ levels in CAD5 cells expressing WT and G127V bvPrP. ImageJ was used to quantify western blots in Figs 4C-G of infected CAD5 cells co-expressing WT bvPrP and G127V bvPrP after exposure to the following prion strains: mouse 22L**(A)**; mouse RML or ME7**(B)**; **(C)** hamster 263K, 139H, DY, or HY; cervid (Deer or Elk) **(D)**; recombinant (Rec A, B, or C) **(E)**. For all statistical analyses, PrP^Sc^ bands were normalized to the PrP^C^/actin ratios of uninfected cells. Bars show the normalized PrP^Sc^ signal relative to chronically infected WT+WT cells set to 100%. p<0.0001 = ****; p<0.001 = ***; p <0.01 = **; ns= not significant.

**Fig. S4:**
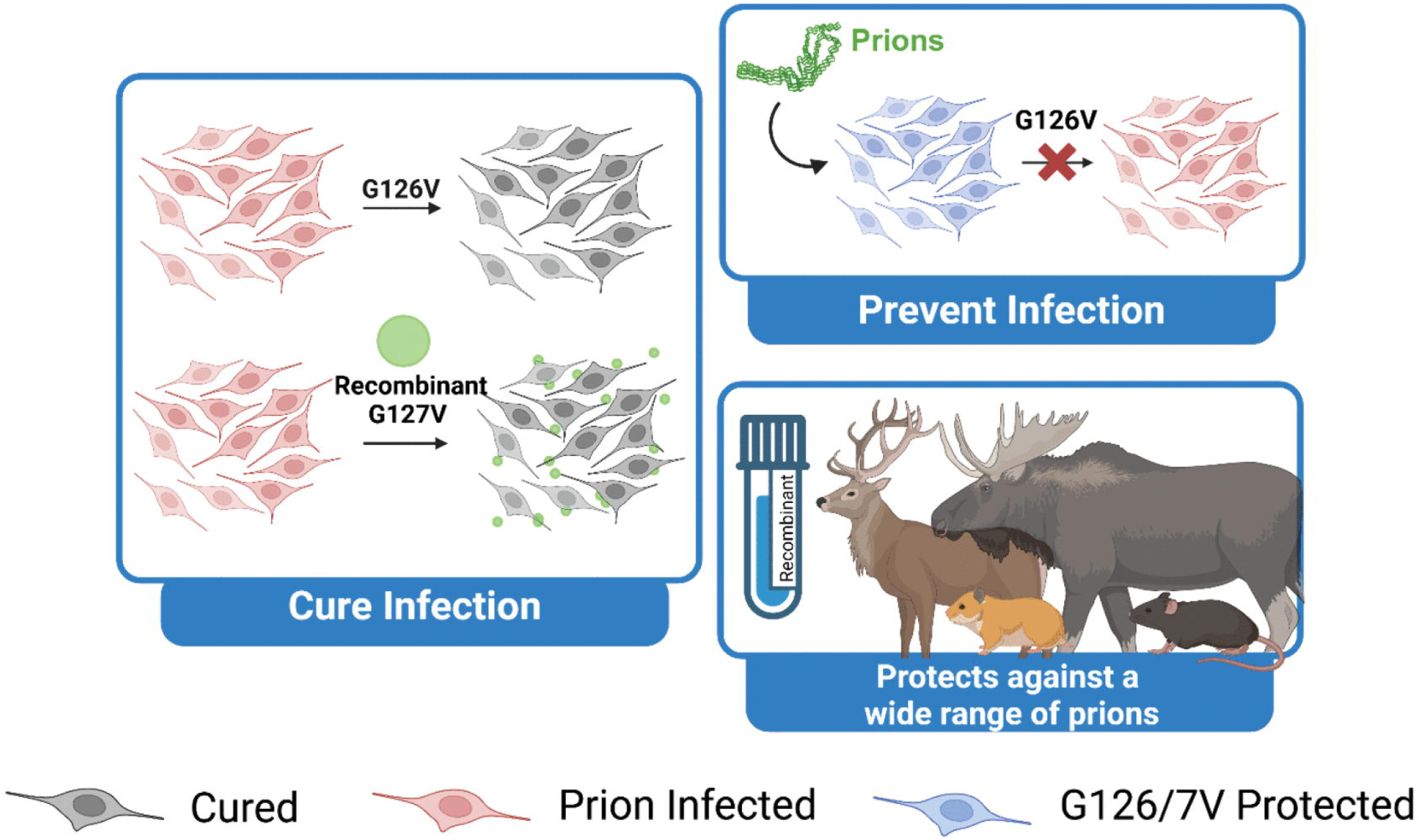
Graphical abstract. Created in BioRender. Gatdula, J. (2026) https://BioRender.com/9y1tsdd

