## Supplemental Material: Raw Western Blots for "Leveraging the dominant-negative effect of the kuru-protective G127V prion protein variant as a novel therapeutic strategy"

### Slide 1
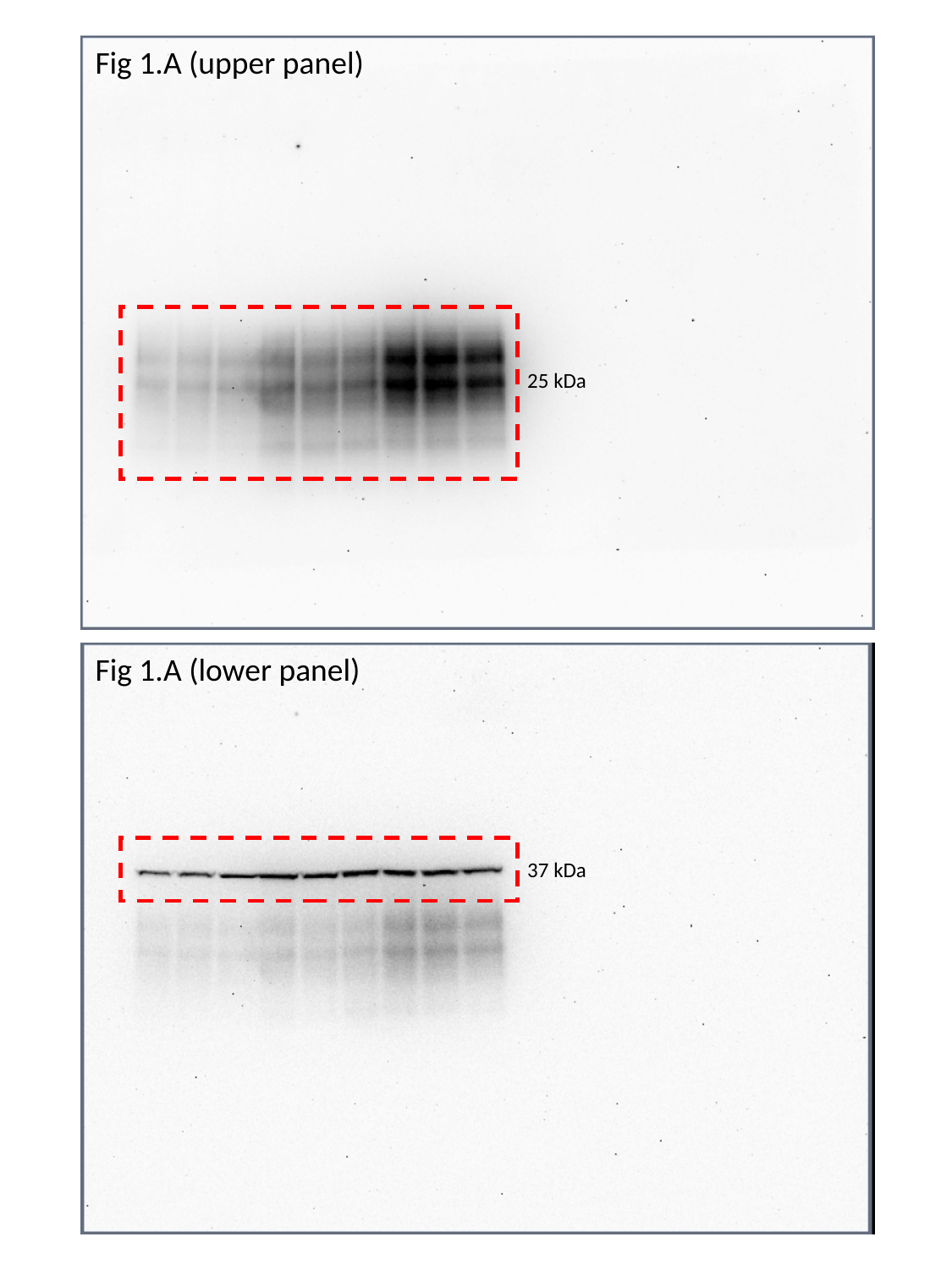

Fig 1.A (upper panel)
25 kDa
Fig 1.A (lower panel)
37 kDa

### Slide 2
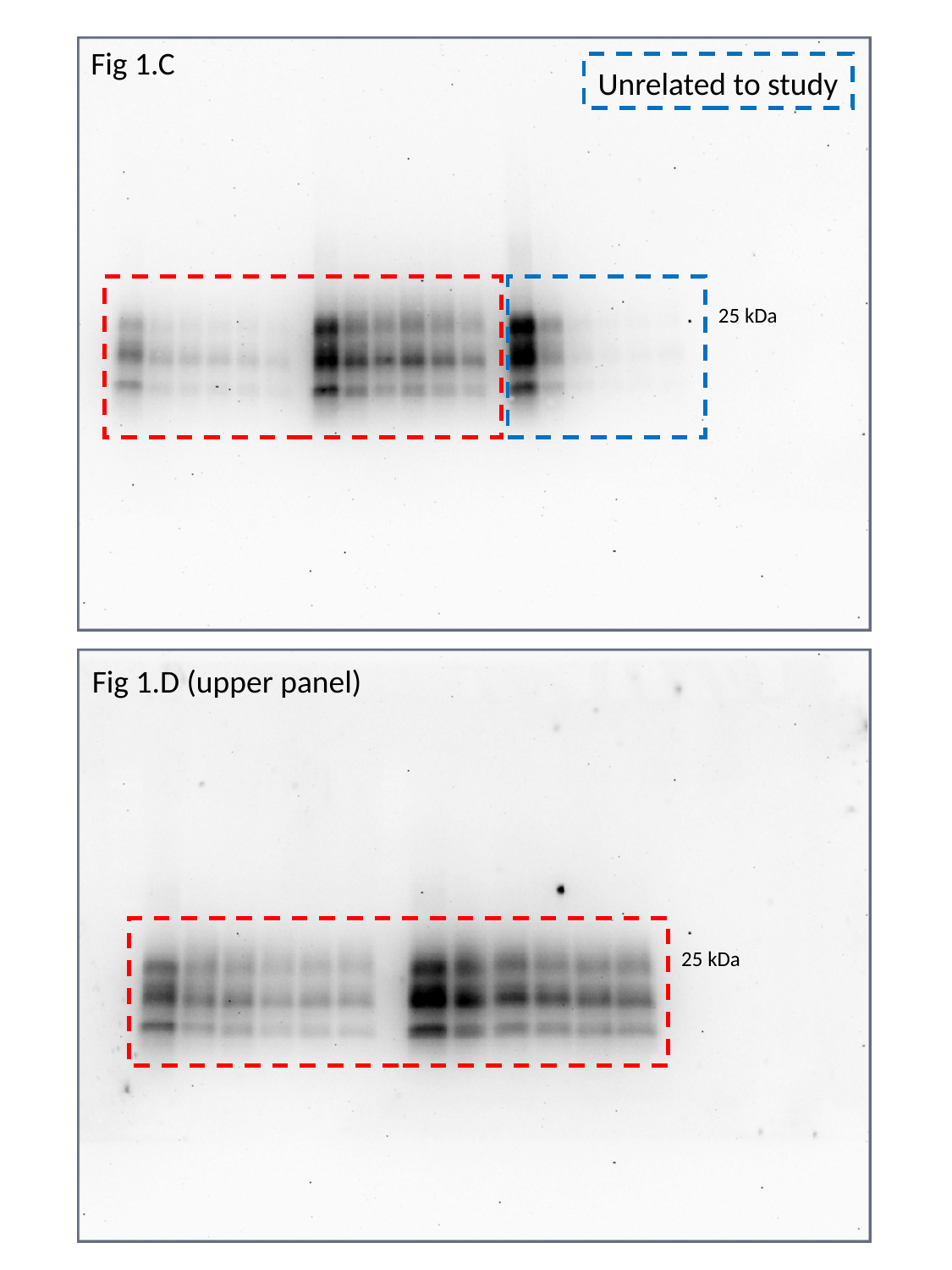

Fig 1.C
Unrelated to study
25 kDa
Fig 1.D (upper panel)
25 kDa

### Slide 3
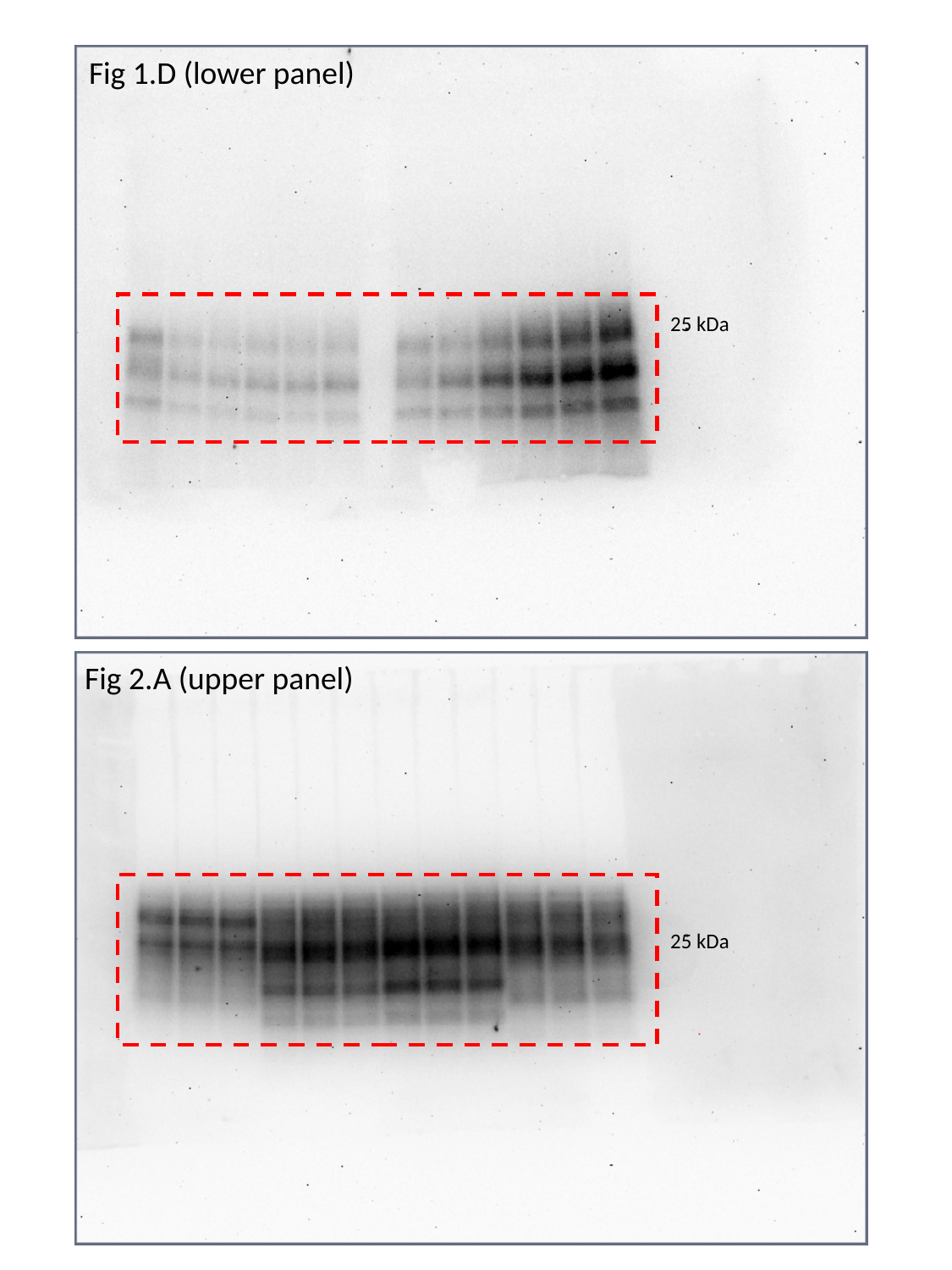

Fig 1.D (lower panel)
25 kDa
Fig 2.A (upper panel)
25 kDa

### Slide 4
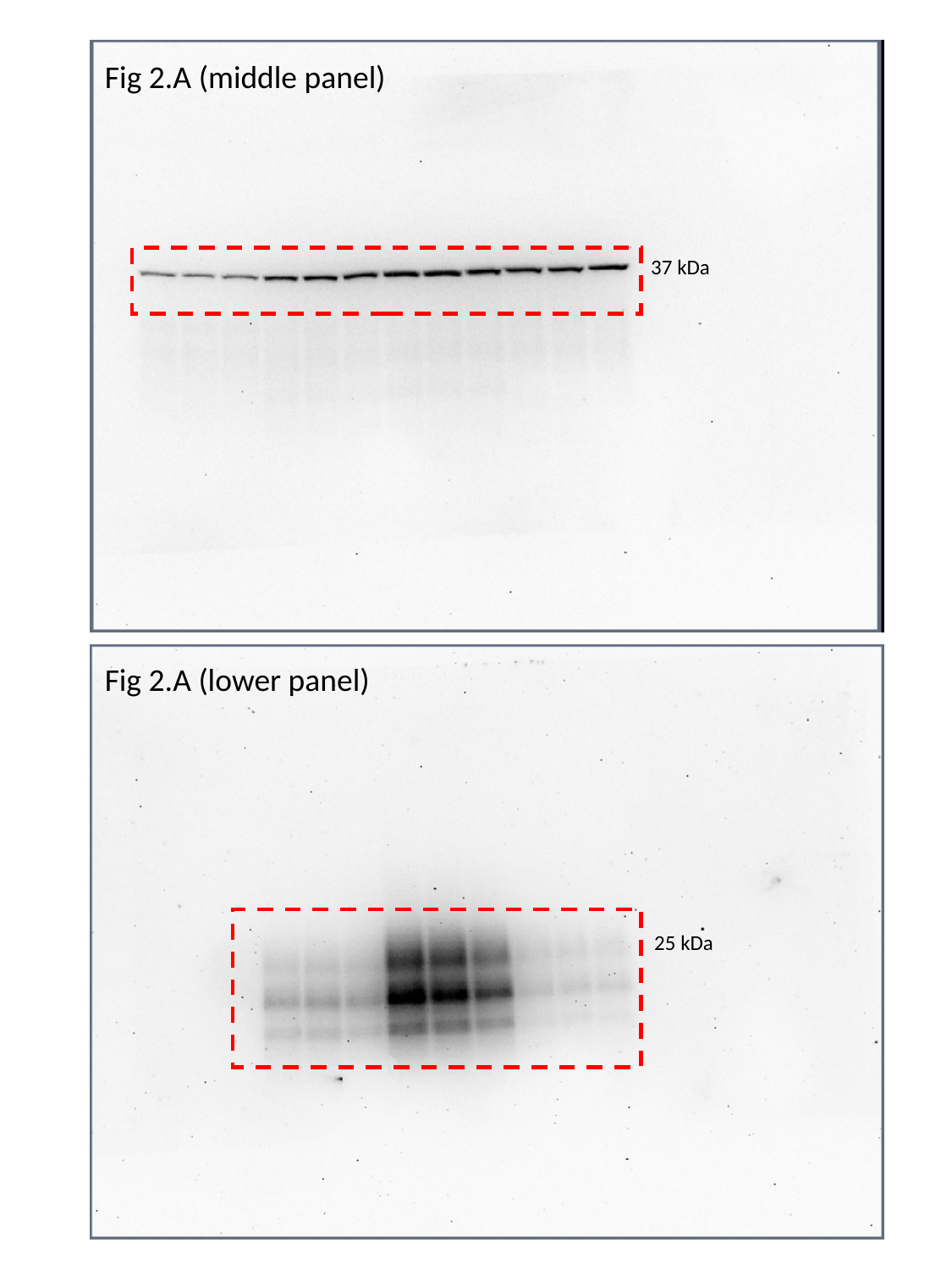

Fig 2.A (middle panel)
37 kDa
Fig 2.A (lower panel)
25 kDa

### Slide 5
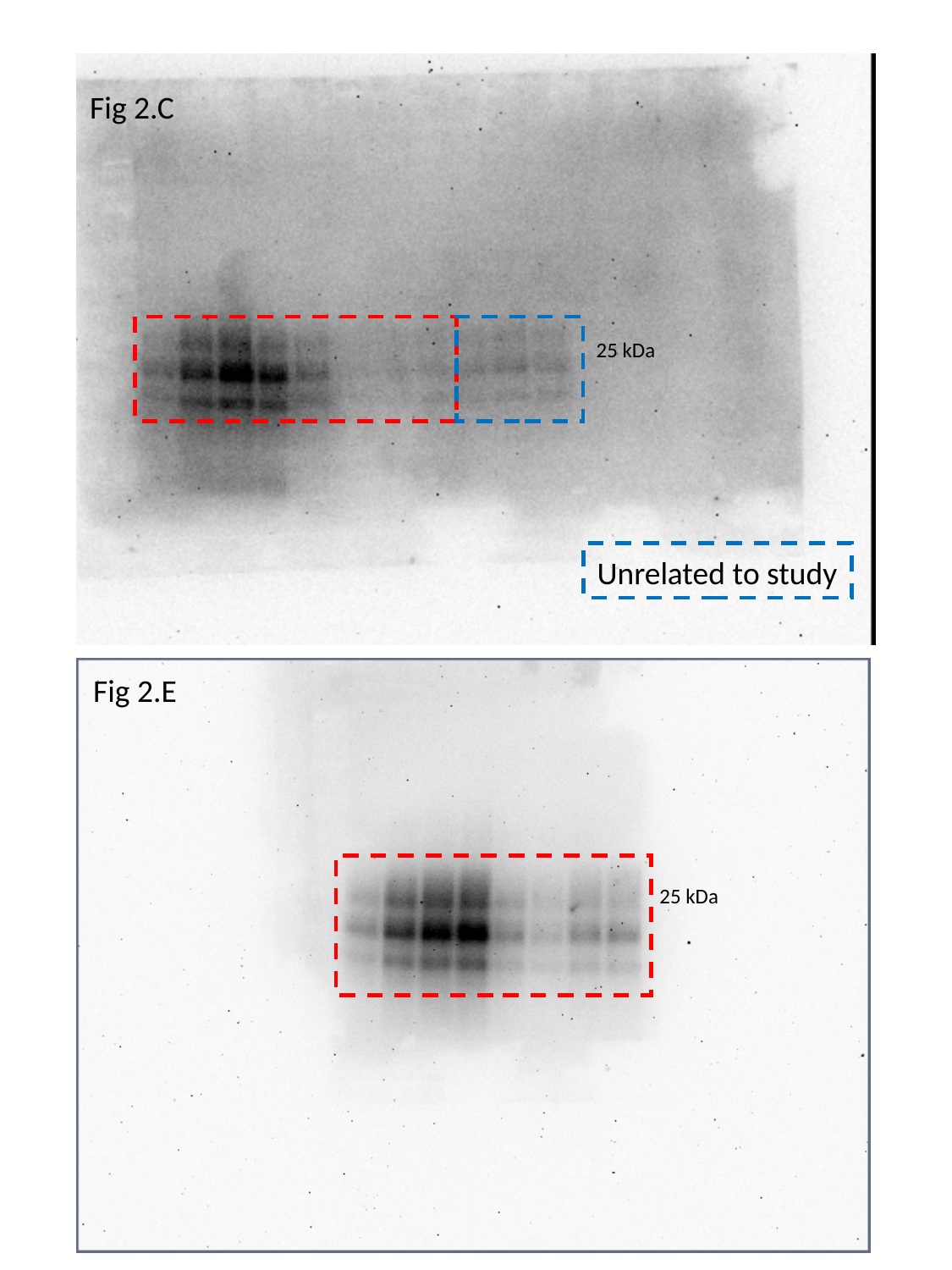

Fig 2.C
25 kDa
Unrelated to study
Fig 2.E
25 kDa

### Slide 6
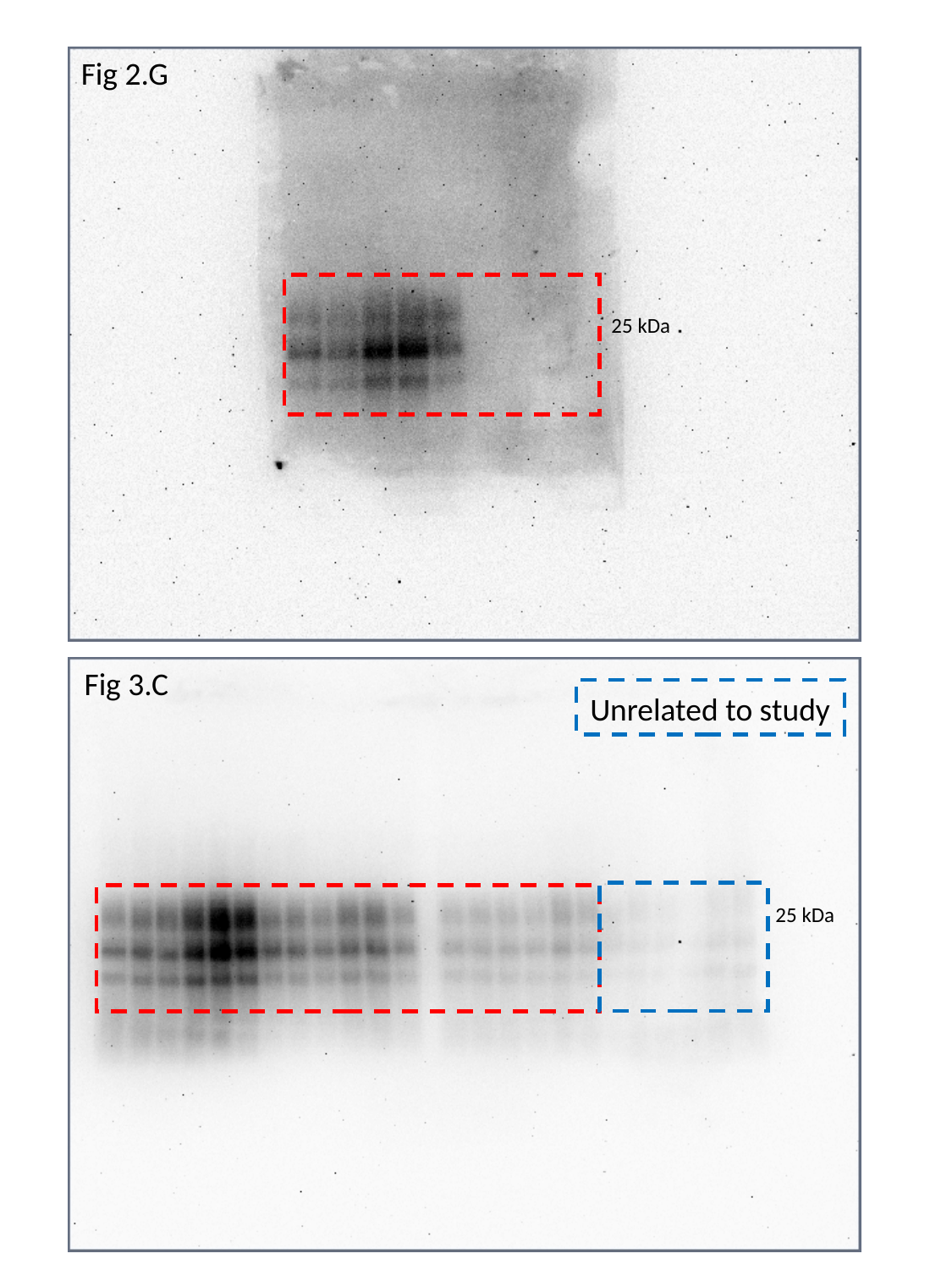

Fig 2.G
25 kDa
Fig 3.C
Unrelated to study
25 kDa

### Slide 7
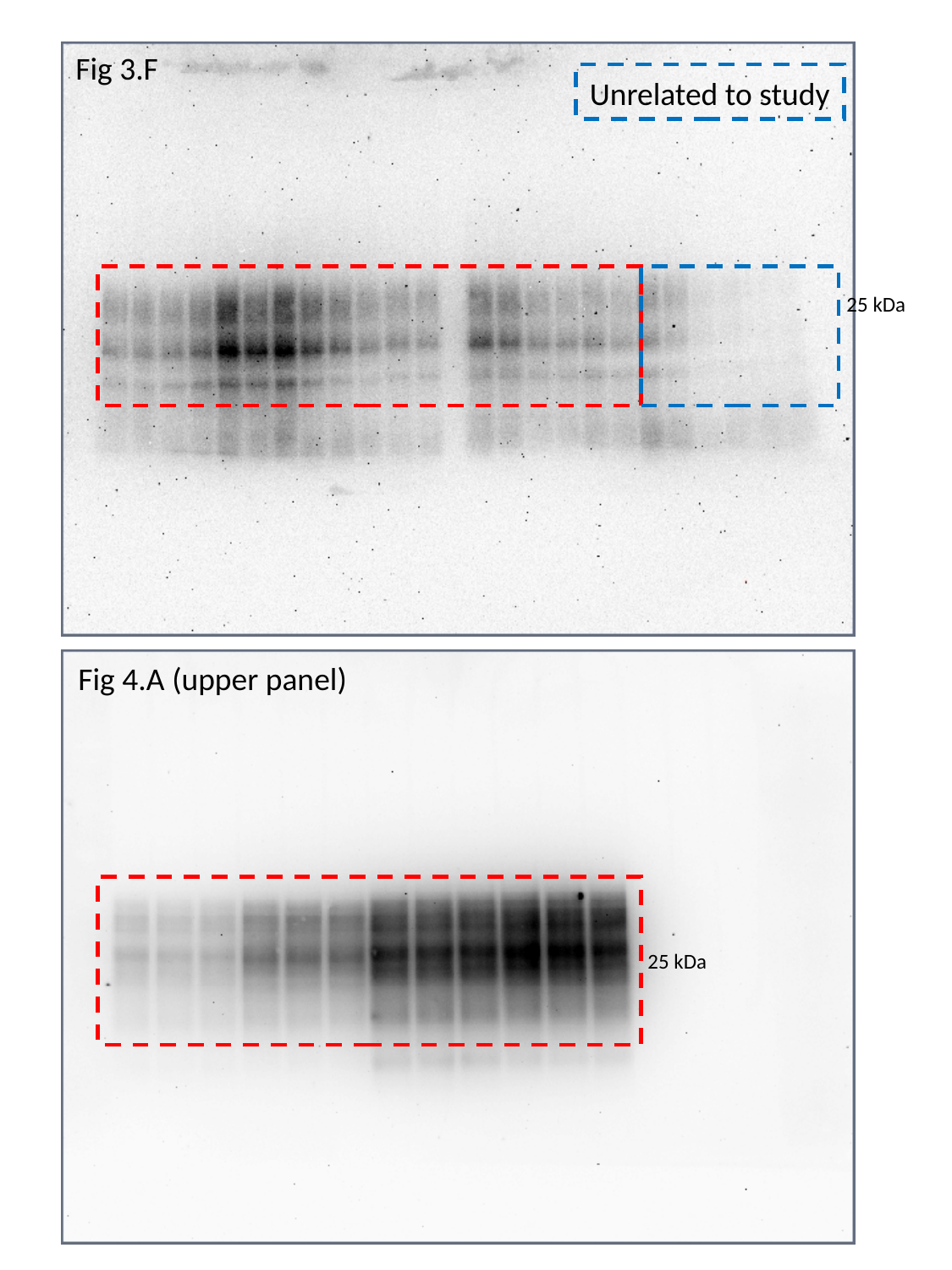

Fig 3.F
Unrelated to study
25 kDa
Fig 4.A (upper panel)
25 kDa

### Slide 8
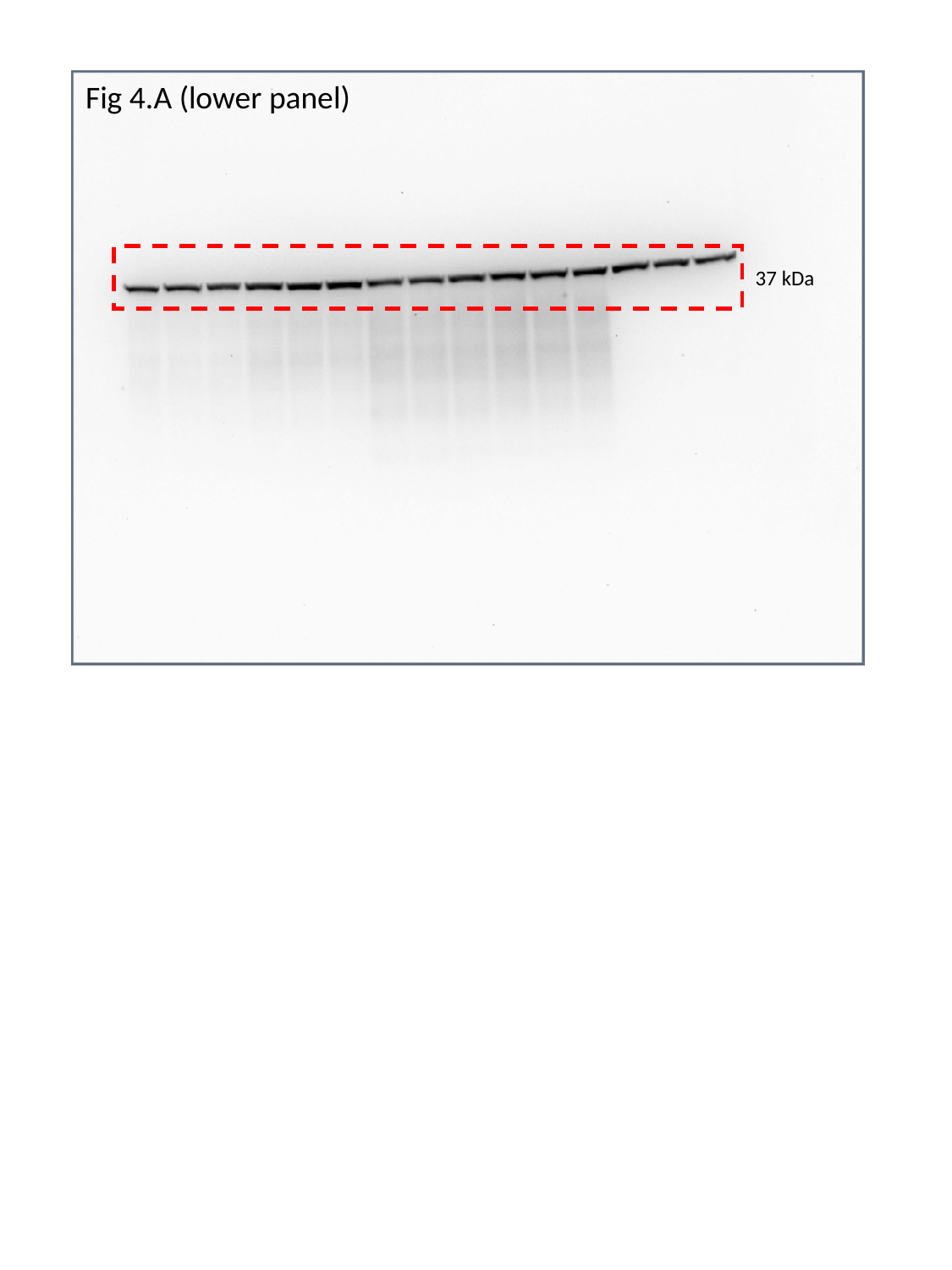

Fig 4.A (lower panel)
37 kDa

### Slide 9
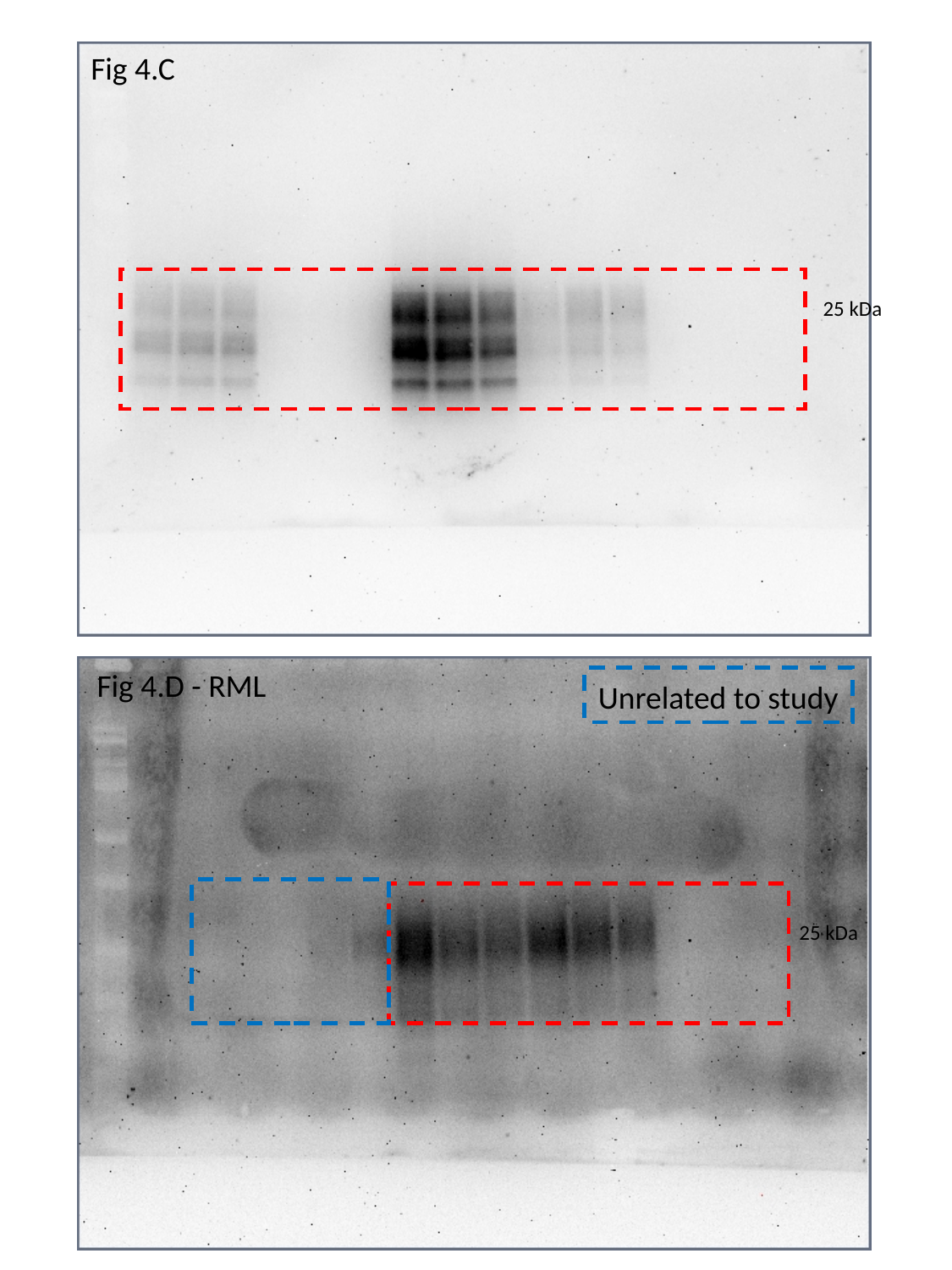

Fig 4.C
25 kDa
Fig 4.D - RML
Unrelated to study
25 kDa

### Slide 10
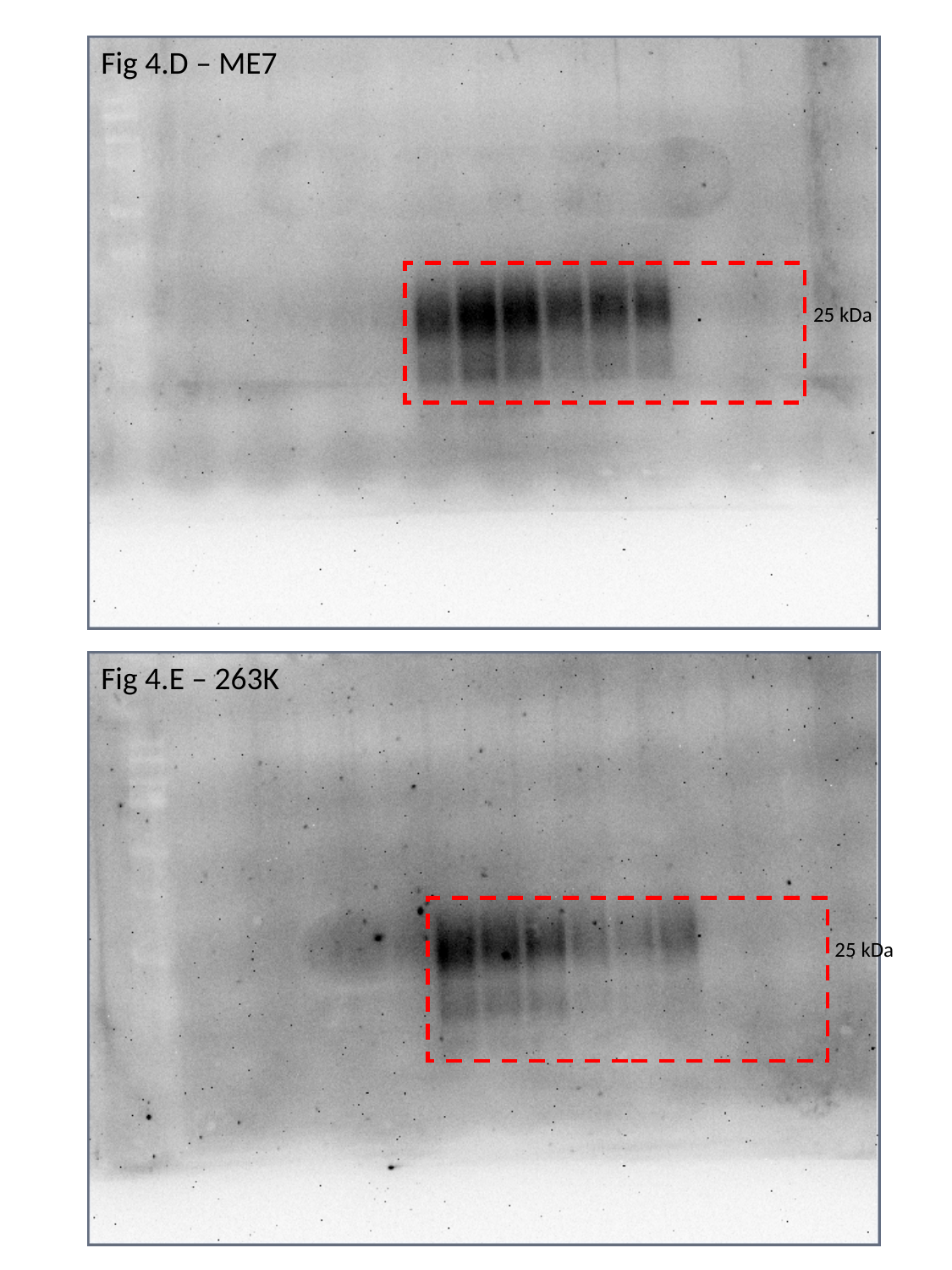

Fig 4.D – ME7
25 kDa
Fig 4.E – 263K
25 kDa

### Slide 11
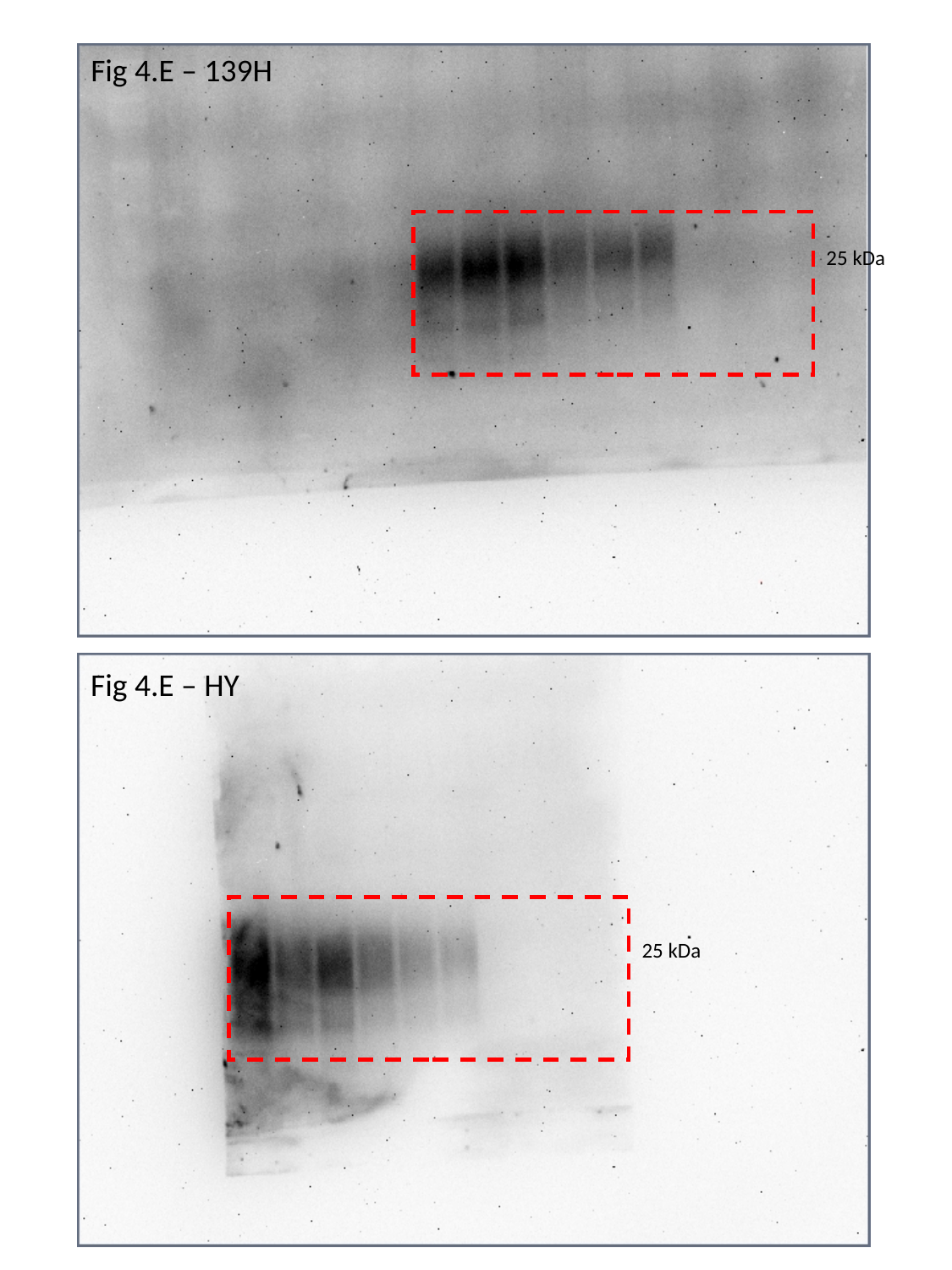

Fig 4.E – 139H
25 kDa
Fig 4.E – HY
25 kDa

### Slide 12
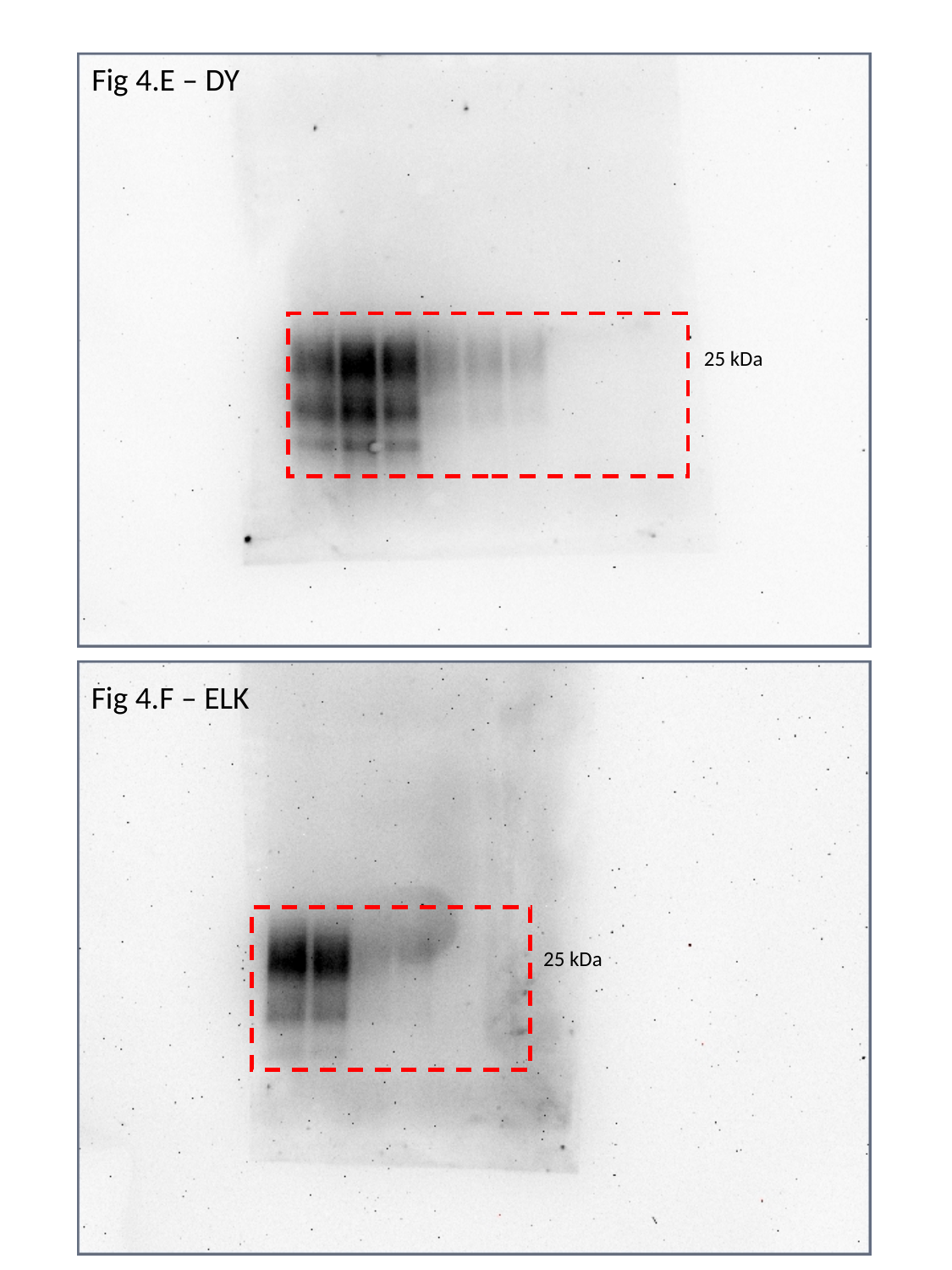

Fig 4.E – DY
25 kDa
Fig 4.F – ELK
25 kDa

### Slide 13
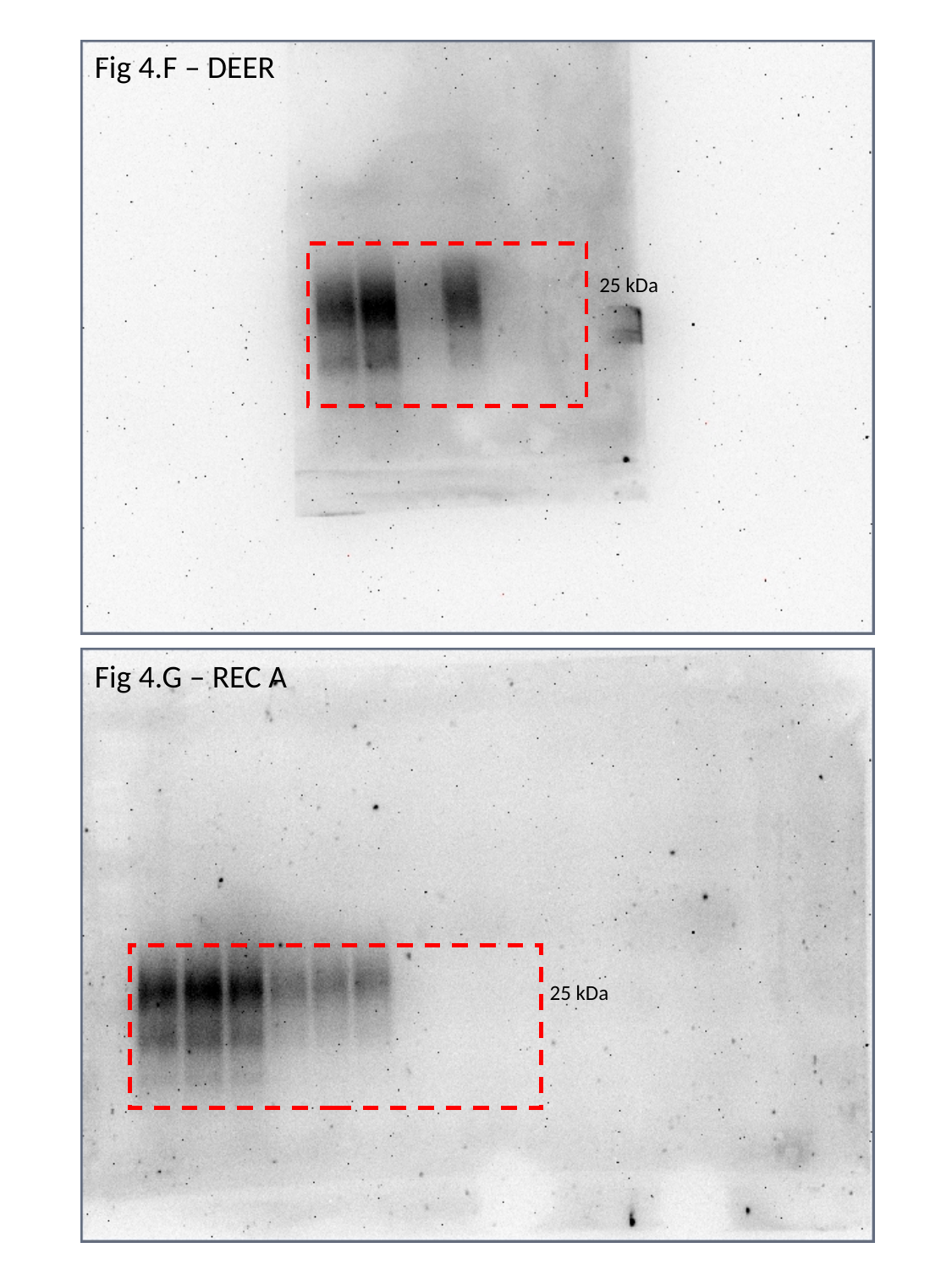

Fig 4.F – DEER
25 kDa
Fig 4.G – REC A
25 kDa

### Slide 14
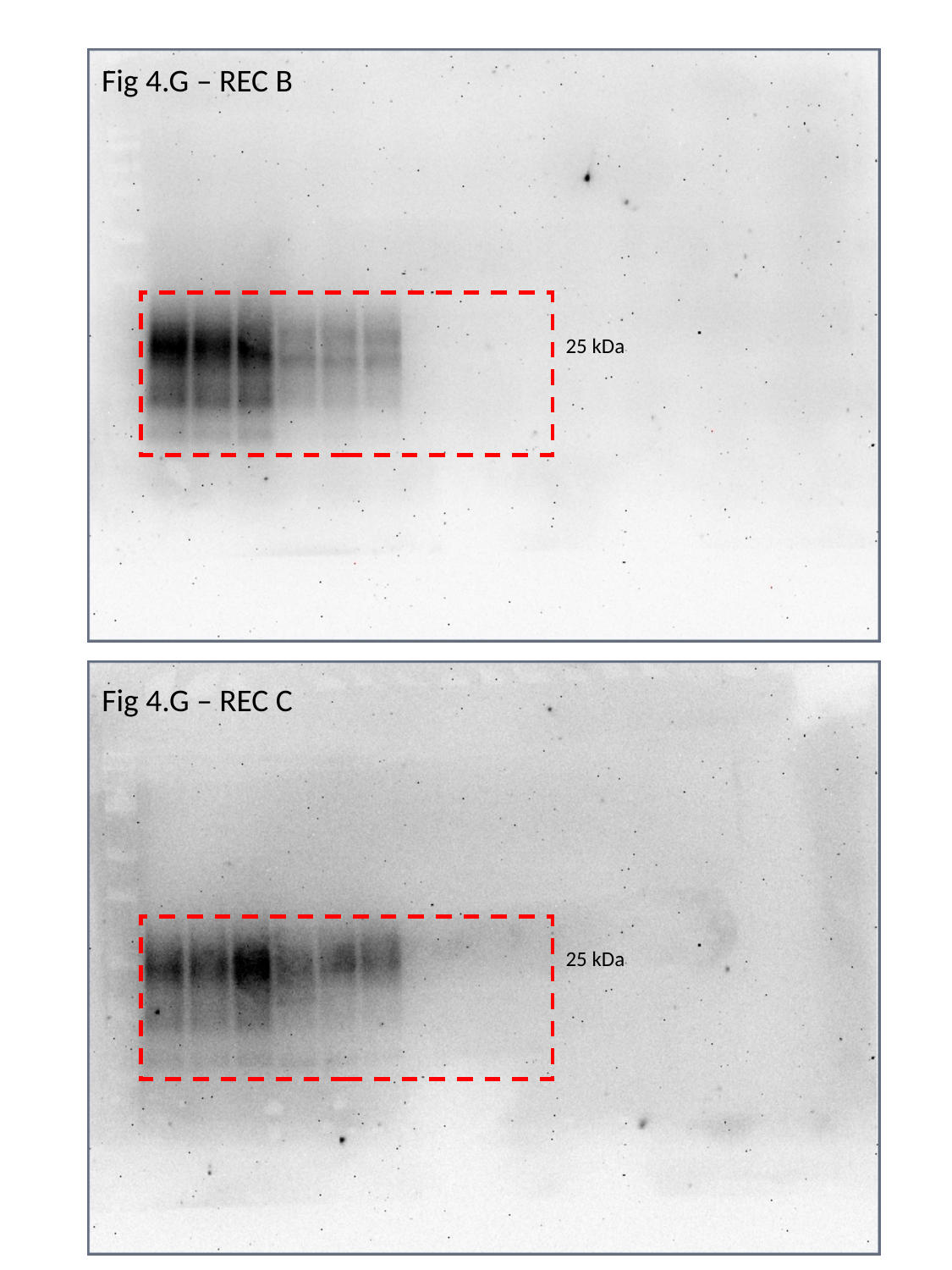

Fig 4.G – REC B
25 kDa
Fig 4.G – REC C
25 kDa

### Slide 15
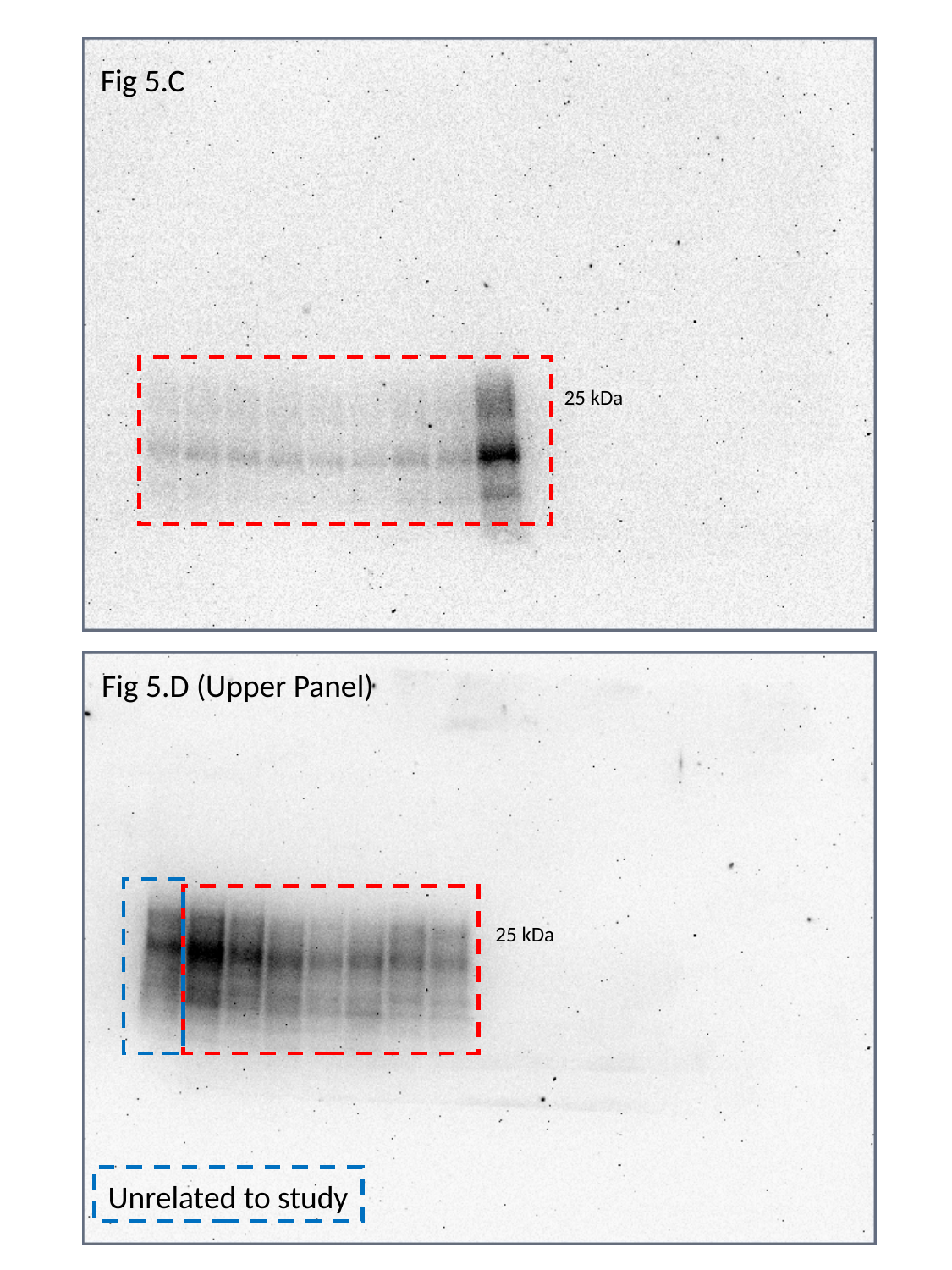

Fig 5.C
25 kDa
Fig 5.D (Upper Panel)
25 kDa
Unrelated to study

### Slide 16
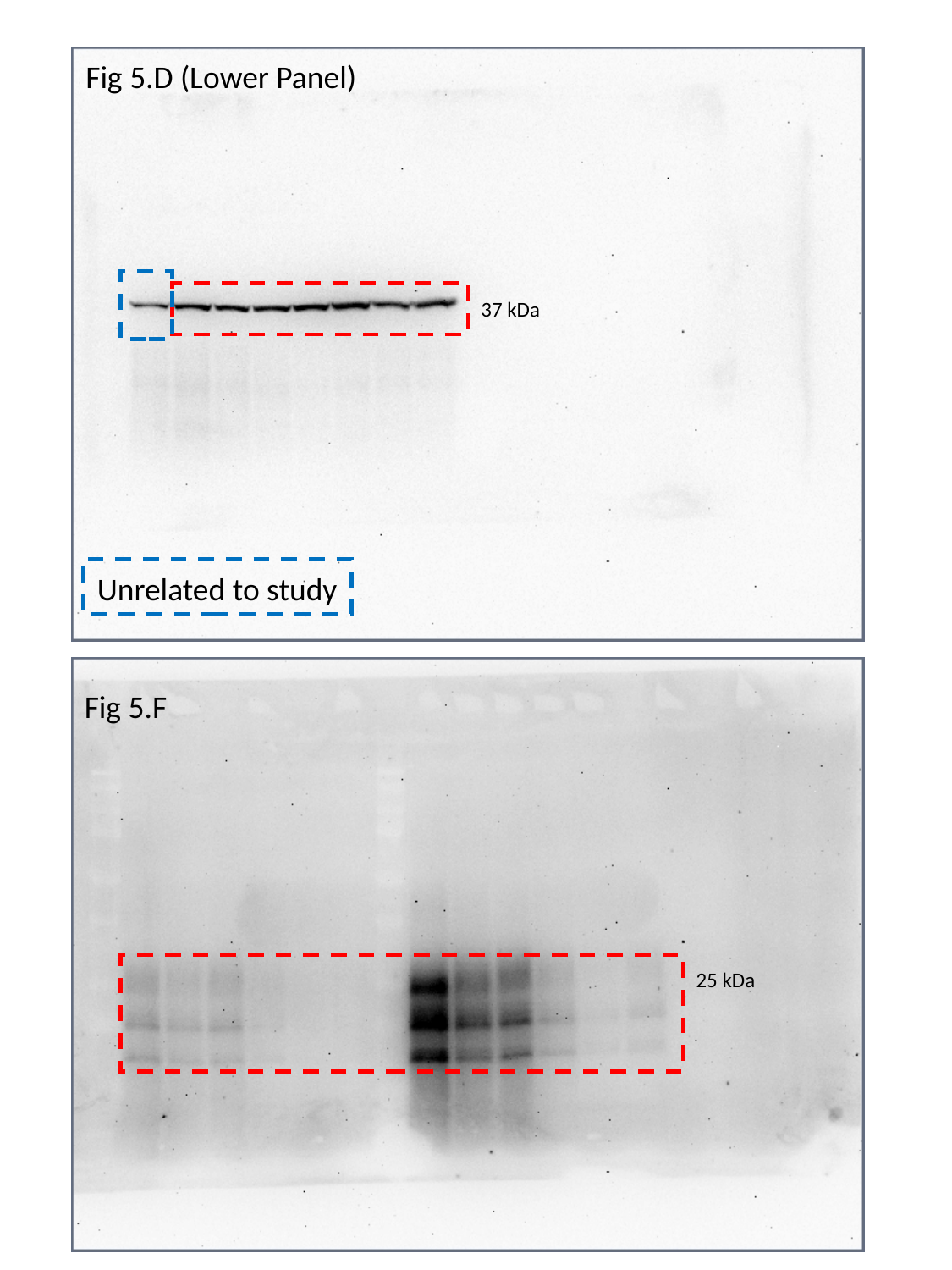

Fig 5.D (Lower Panel)
37 kDa
Unrelated to study
Fig 5.F
25 kDa

### Slide 17
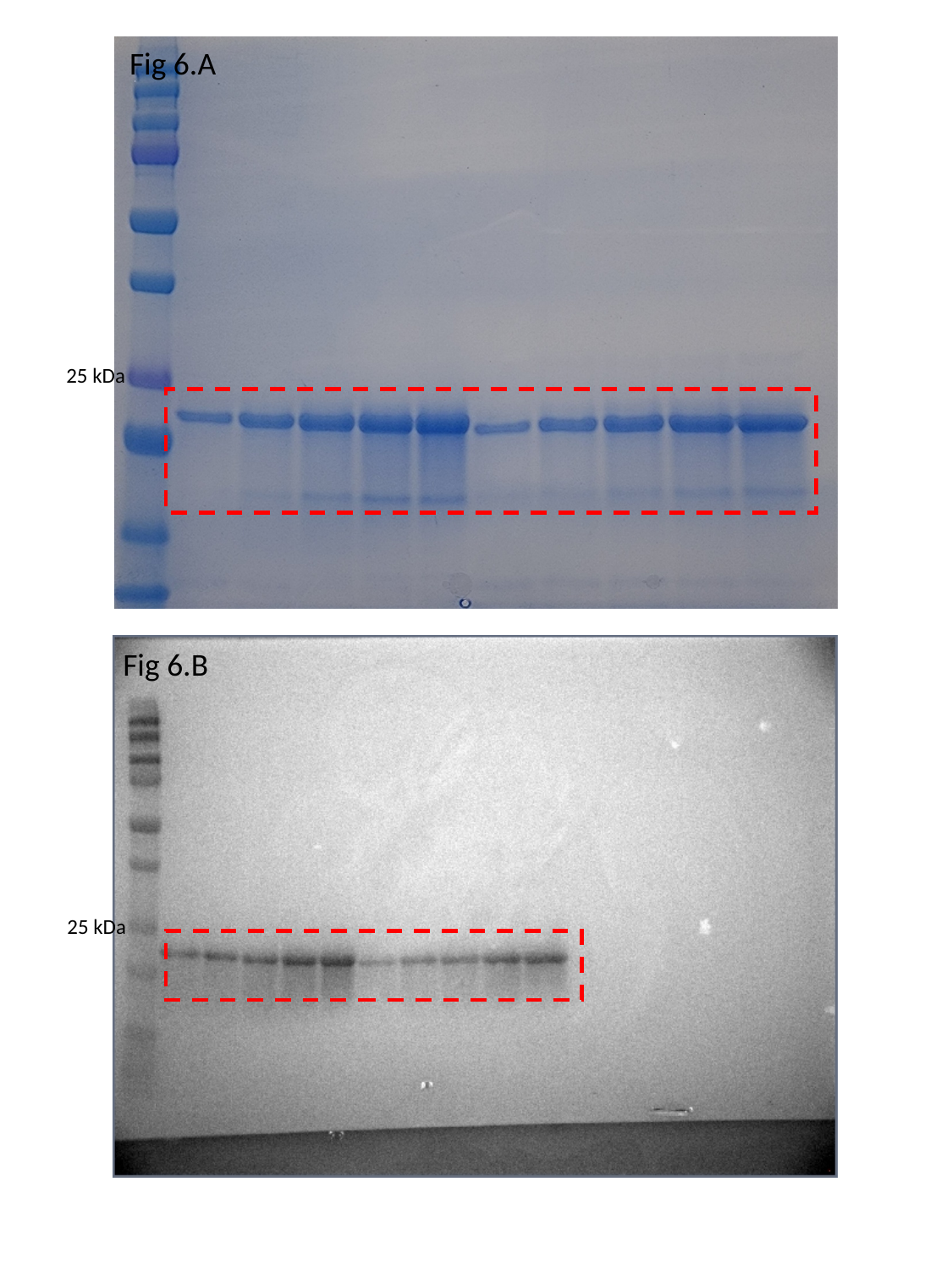

Fig 6.A
25 kDa
Fig 6.B
25 kDa

### Slide 18
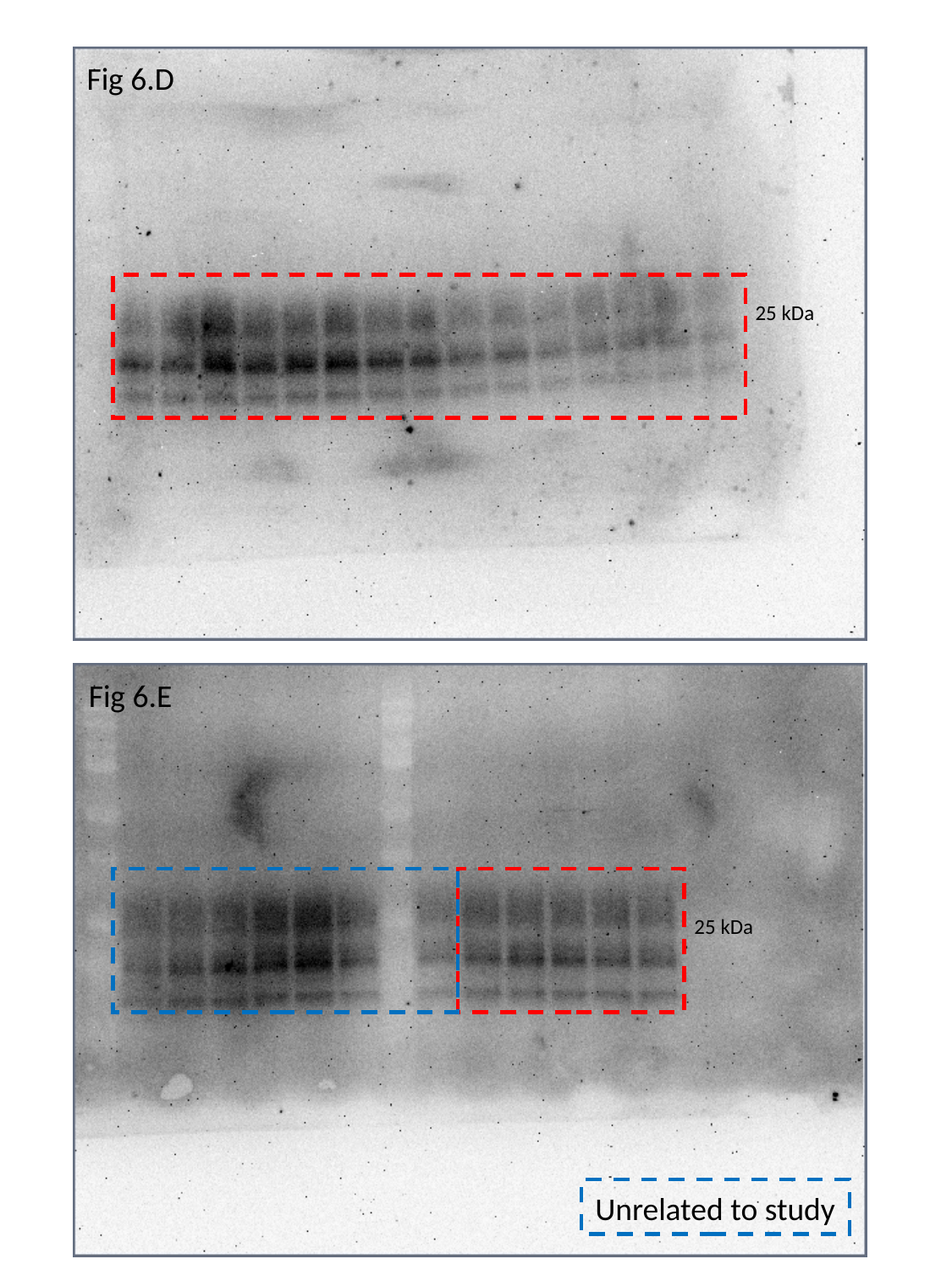

Fig 6.D
25 kDa
Fig 6.E
25 kDa
Unrelated to study

### Slide 19
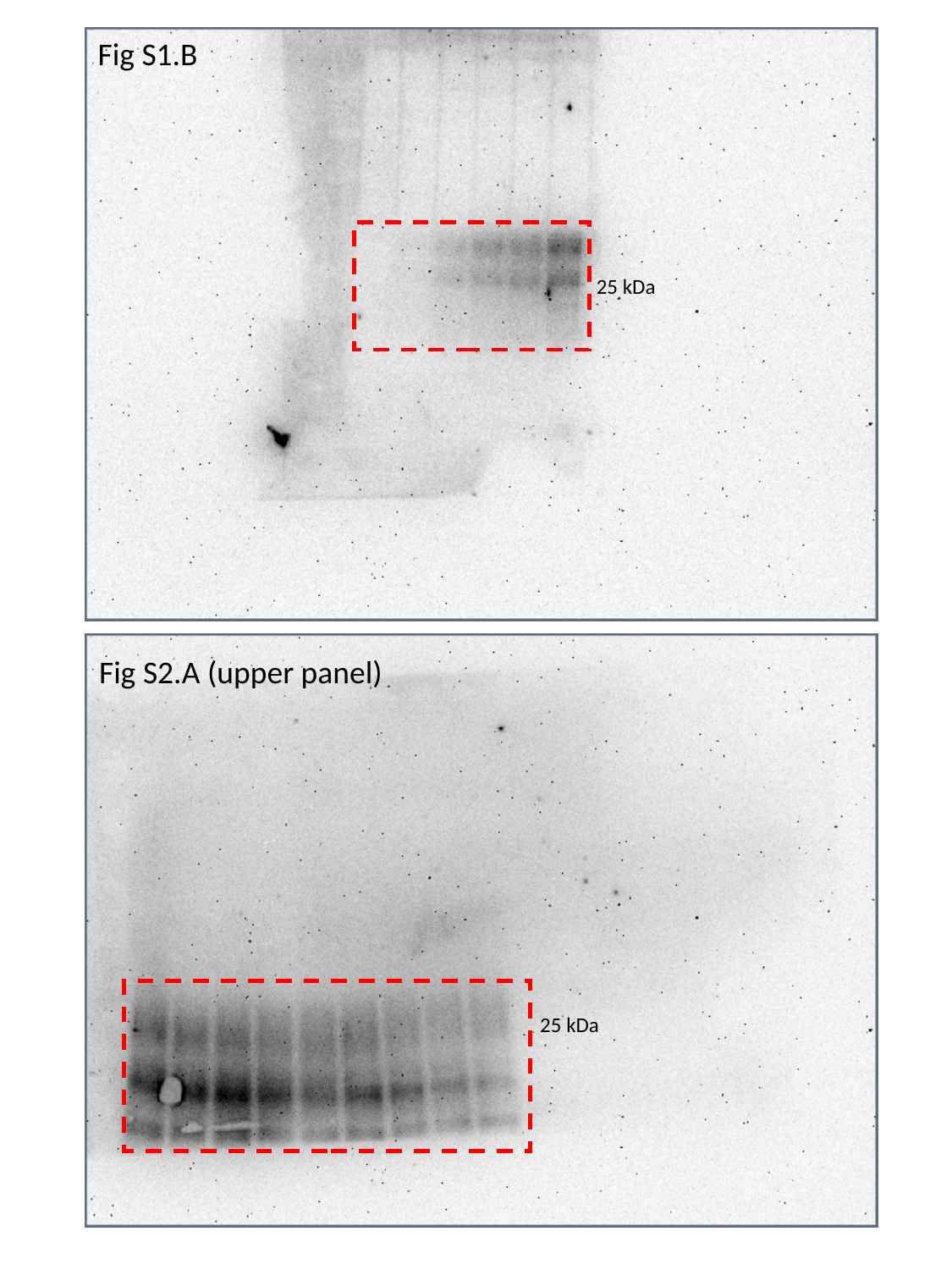

Fig S1.B
25 kDa
Fig S2.A (upper panel)
25 kDa

### Slide 20
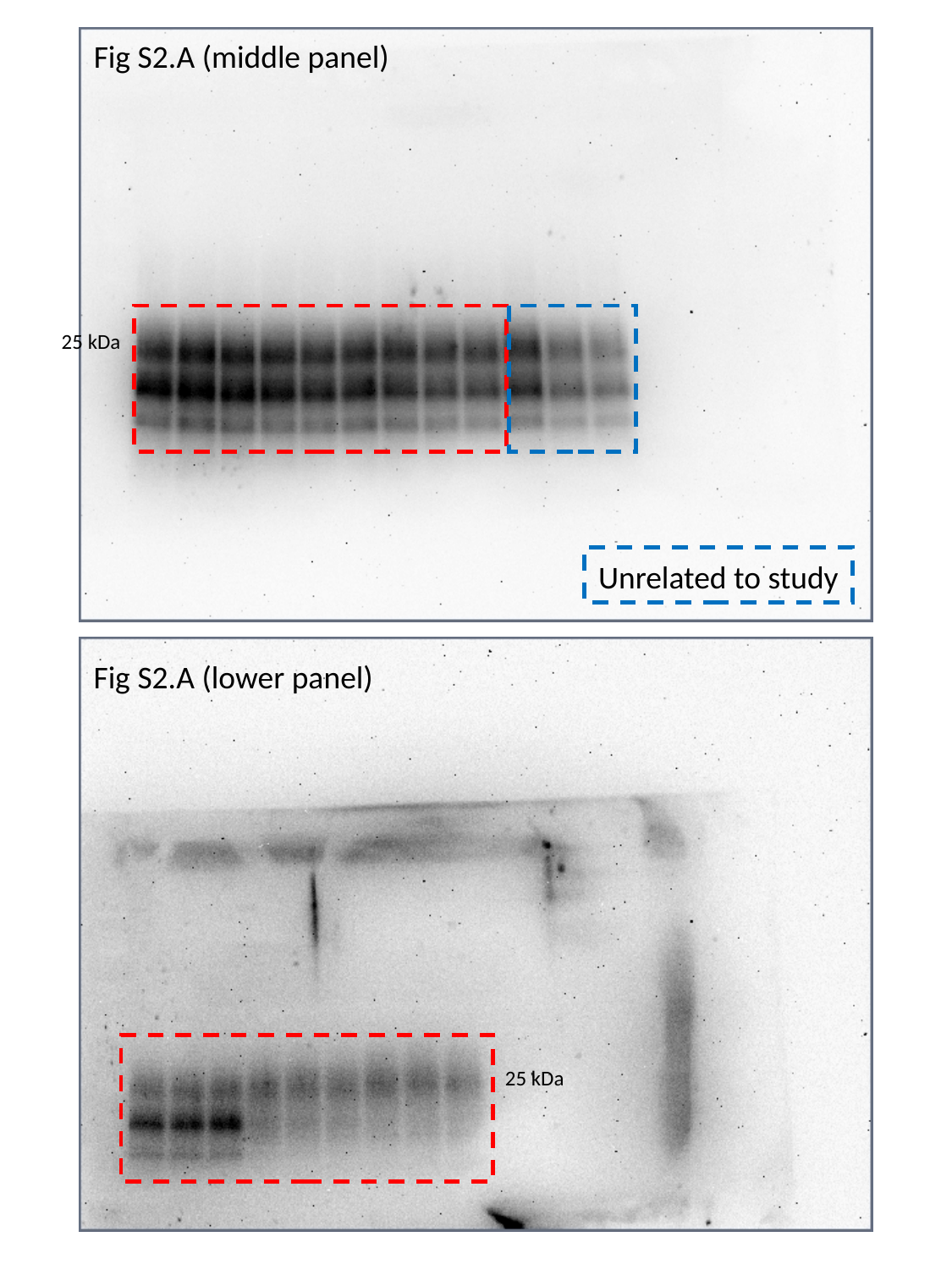

Fig S2.A (middle panel)
25 kDa
Unrelated to study
Fig S2.A (lower panel)
25 kDa
